# ARGLU1 is a Glucocorticoid Receptor Coactivator and Splicing Modulator Important in Stress Hormone Signaling and Brain Development

**DOI:** 10.1101/069161

**Authors:** Lilia Magomedova, Jens Tiefenbach, Emma Zilberman, Veronique Voisin, Melanie Robitaille, Serge Gueroussov, Manuel Irimia, Debashish Ray, Rucha Patel, ChangJiang Xu, Pancharatnam Jeyasuria, Gary D. Bader, Timothy R. Hughes, Henry Krause, Benjamin J. Blencowe, Stephane Angers, Carolyn L. Cummins

## Abstract

Prolonged exposure to glucocorticoid stress hormones precipitates mood and cognitive disorders. We identified arginine and glutamate rich 1 (ARGLU1) in a screen for new modulators of glucocorticoid signaling in the CNS. Biochemical studies found that the glutamate rich C-terminus coactivates the glucocorticoid receptor (GR) and the arginine rich N-terminus interacts with splicing factors and RNA. RNA-seq of neuronal cells ±siARGLU1found significant changes in the expression and alternative splicing of distinct genes involved in neurogenesis. Loss of ARGLU1 was embryonic lethal in mice, and knockdown in zebrafish caused neurodevelopmental and heart defects. Treatment with dexamethasone, a GR activator, also induced changes in the pattern of alternatively spliced genes, highlighting an underappreciated global mechanism of glucocorticoid action in neuronal cells. Thus, in addition to its basal role, ARGLU1 links glucocorticoid-mediated transcription and alternative splicing in neural cells, providing new avenues from which to investigate the molecular underpinnings of cognitive stress disorders.

## Highlights

- ARGLU1 is a new GR coactivator that is nuclear localized and highly expressed in the CNS.
- Dexamethasone, a GR ligand, induces alternative splicing changes in neural cells that are ARGLU1-dependent.
- ARGLU1 impacts two layers of gene regulation (transcription and alternative splicing) on largely mutually exclusive genes.
- Loss of ARGLU1 is embryonic lethal in mice and knockdown in zebrafish causes heart and brain defects.

## Short Summary

Stress hormones, such as cortisol, significantly alter developing and adult neurons by signaling through the glucocorticoid receptor. We identified ARGLU1 as a glucocorticoid-receptor coactivator and RNA binding protein implicated in the regulation of both transcription and alternative splicing in neuronal cells. Glucocorticoid treatment caused widespread changes in alternative splicing that were abrogated in the absence of ARGLU1. Our data support a critical role for ARGLU1 in both basal and glucocorticoid-mediated alternative splicing and transcription, processes important for neuronal development.

The glucocorticoid receptor (GR) plays a fundamental role in coordinating the transcriptional response to stress hormones, such as cortisol, and is essential for organismal development, glucose homeostasis and immune function (Kadmiel and Cidlowski, 2013). Chronic stress, or administration of glucocorticoid drugs, is known to impair working memory and precipitate the onset of neuropsychiatric disorders (Judd et al., 2014; Lupien et al., 2007). In the brain, GR is highly expressed in the hippocampus where it has been shown to play a central role in modulating the proliferation and differentiation of neural stem and progenitor cells (Fitzsimons et al., 2016; Fitzsimons et al., 2013; Mahfouz et al., 2016).

GR and other members of the nuclear receptor superfamily share highly conserved domains including the zinc-finger DNA-binding domain (DBD) and a carboxy terminal ligand-binding domain (LBD) attached to a ligand dependent activation function domain (AF2). In the absence of ligand, GR is complexed to chaperone proteins in the cytosol that dissociate upon ligand binding and unmask a nuclear localization signal. GR then translocates to the nucleus where it regulates gene expression by binding to GR response DNA elements or to other transcription factors.

Ligand-bound GR is known to interact with members of the p160 coactivator family including steroid receptor coactivator 1 (SRC1) and glucocorticoid receptor interacting protein GRIP1 (TIF2). SRC1 and TIF2 can recruit histone acetyltransferase enzymes, resulting in changes in chromatin structure (Chen et al., 1997; Huang and Cheng, 2004; Spencer et al., 1997). Interactions between coactivators and other transcription factors and regulatory proteins help to recruit RNA polymerase II and initiate gene transcription (Robyr et al., 2000). Coregulators play roles in every step of transcription including chromatin remodeling, initiation, elongation, and termination (Lonard et al., 2007). Some coregulators have also been implicated in the regulation of RNA splicing (Auboeuf et al., 2007; Auboeuf et al., 2004; Auboeuf et al., 2002; Huang et al., 2012; Sun et al., 2007; Zhang et al., 2003). While numerous GR coregulators have been identified, their effects on stress-induced corticosteroid signaling in the brain remain largely unexplored. To identify new biological mediators of GR function in the central nervous system (CNS), we performed a high-throughput expression cloning screen examining GR transcriptional activity in response to stress hormones.

Herein, we report that arginine and glutamate rich 1 (ARGLU1) is a highly evolutionarily conserved GR coactivator and RNA splicing modulator. We show in neuronal cells that glucocorticoid signaling, through dexamethasone treatment, not only affects transcription but significantly changes the alternative splicing landscape in an ARGLU1-dependent manner. These functions were previously unknown as ARGLU1 had only been shown to interact with the mediator complex MED1 and affect estrogen receptor signaling (Zhang et al., 2011). We show that ARGLU1 is highly expressed in the CNS and loss of ARGLU1 is embryonic lethal in mice. Our data support a model in which ARGLU1 uses its distinct domains to control mRNA on two levels: by changing gene expression and alternative splicing in selected pathways such as histone chromatin organization and neurogenesis. The widespread glucocorticoid (GC)-mediated changes in alternative splicing found in neuronal cells highlight a new avenue from which to explore the molecular mechanisms by which stress hormones impact mood, memory and cognition.

## Results

### ARGLU1 is identified as a transcriptional coactivator of GR from a cDNA expression cloning screen

In an effort to identify novel proteins influencing stress hormone signaling, an expression cloning assay measuring GR activity was optimized in HEK293 cells and screened in the presence of cDNA pools from a normalized human brain cDNA library. The GC-responsive reporter system consisted of a plasmid encoding the fusion protein of human GR ligand binding domain linked to the yeast GAL4 DNA binding domain (GAL4-GR), and a luciferase reporter plasmid driven by four GAL4 upstream activating sequences (UAS-luc). Cortisol, an endogenous GR ligand and stress hormone, was used at a sub-saturating dose of 300 nM.

The maximum number of cDNAs that could be readily screened per well in a high-throughput format in a 96-well plate was determined by dilution experiments of positive controls. A 2-fold increase or decrease in luciferase activity with co-transfection of the known GR coactivator, TIF2, and corepressor, RIP140, respectively, was detectable when diluted 50-fold with an empty plasmid, indicating that a cDNA pool size of ∼50 could be effectively screened in our assay. By serial dilution of the cDNA library after electroporation, it was estimated that the library contained 1.6 × 10^9^ independent transformants. A total of ∼105,600 cDNAs in 2112 pools were screened in a 96-well format (22 plates total containing 50 clones/well). Hits were considered positive if they yielded a luciferase signal greater than 2-fold compared to control transfected cells. Subsequent rounds of sib selection were performed screening at 12 clones/well and 1 clone/well (Figure 1A). Following up one of the hits led to the identification of a cDNA encoding the arginine and glutamate rich 1 (ARGLU1) protein (Figure 1B).

**Figure 1:**
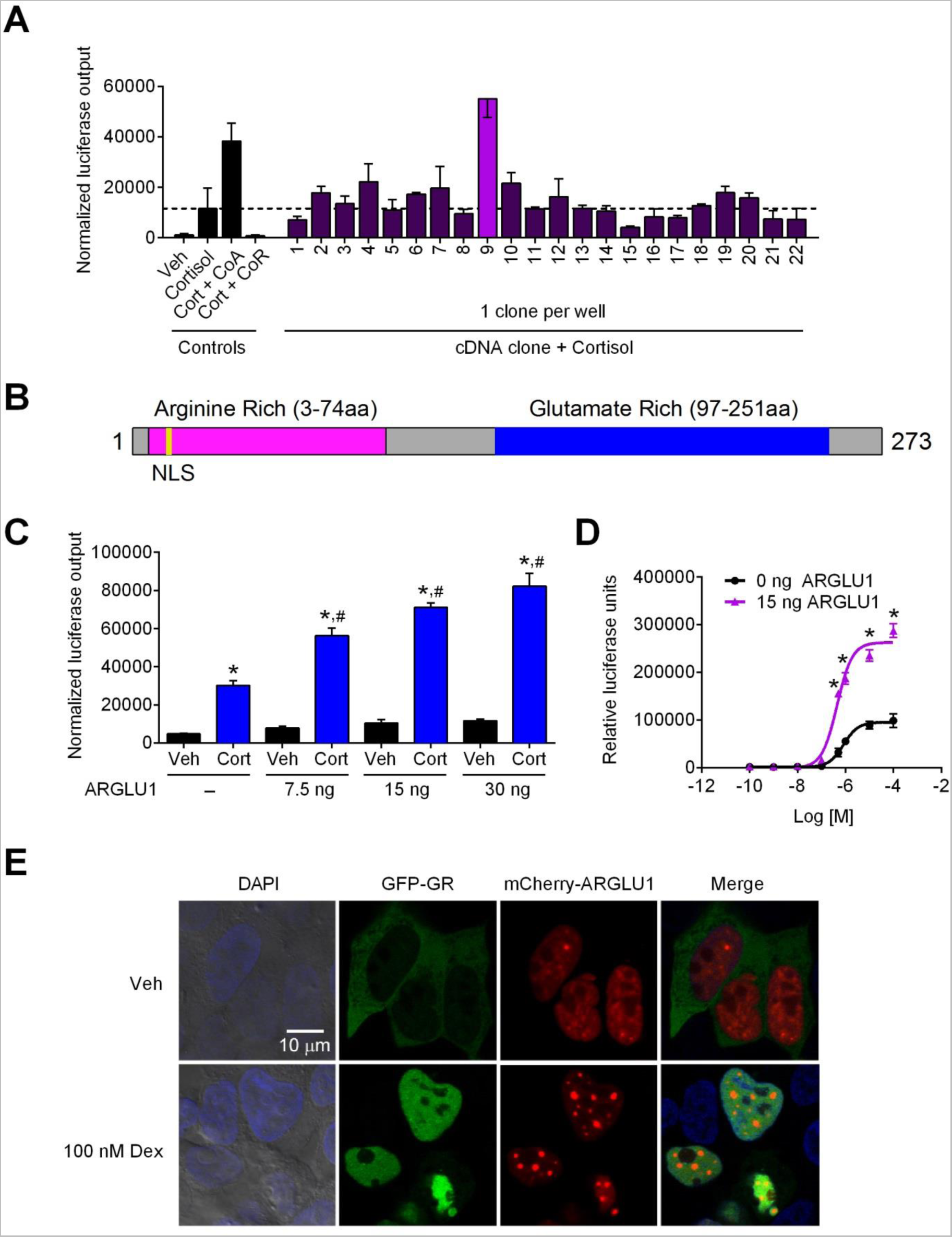
A screen for new modulators of GR activity identified ARGLU1, a nuclear protein from a human brain cDNA library, as a GR coactivator. (A) cDNA pools from a human brain library were co-transfected into HEK293 cells with GAL4-GR, UAS-luciferase and β-galactosidase. Cells were treated with 300 nM cortisol (Cort) and the luciferase signal was compared to no library control. The coactivator (CoA) TIF2 and corepressor (CoR) RIP140 were used as positive controls. The positive hit shown in light purple (>2-fold change) was first screened at 50 clones per well, then 12 clones per well, and finally 1 clone per well (A). The clone in light purple was identified as ARGLU1. (B) Schematic of ARGLU1 domains. NLS, nuclear localization sequence. (C) HEK293 cells transfected with the GAL4-GR/UAS-luc reporter with increasing amounts of CMX-ARGLU1 and 300 nM cortisol. (D) GAL4-GR/UAS-luc transfection with a constant amount of CMX-ARGLU1 (15 ng/well) and increasing concentrations of cortisol. (A, C, D) Data represent the mean ± SD, (C) ANOVA followed by Newman-Keuls, *p < 0.05 Cort vs respective Veh; #p < 0.05 vs 0 ng. (D) Student's t-test *p < 0.05 vs 0 ng ARGLU1. (E) GFP-GR (green) and mCherry-ARGLU1 (red) were co-transfected into HEK293 cells and treated with vehicle (EtOH) or 100 nM Dex for 4 hrs. DAPI (blue) was used to stain the nuclei. See also Figures S1-S2.

To confirm that ARGLU1 was a *bone fide* GR coactivator, we transfected HEK293 cells with increasing amounts of ARGLU1 and observed a dose-dependent increase in GR transcriptional activity (Figure 1C). Likewise, we found that ARGLU1 significantly increased the maximal activation of GAL4-GR when tested in a dose response assay with cortisol, suggesting that ARGLU1 functions as a typical coactivator for this receptor (Figure 1D).

### ARGLU1 is highly evolutionarily conserved and unique among nuclear receptor coactivators

ARGLU1 is a 273 amino acid protein (33 kDa) that is so named because of its two distinct regions: the N-terminus which is rich in positively charged arginine amino acids, and the C-terminus which is composed of glutamate rich amino acids (Figure 1B and Figure S1A). There is high evolutionary conservation between orthologs of ARGLU1 with 99% sequence identity between mouse and human ARGLU1 (Figure S1A). Functional complementation was shown for mouse and zebrafish ARGLU1 when tested in a human GR co-transfection assay (Figure S1B). ARGLU1 is not related to other known NR coregulators and, to date, has only been reported to interact with MED1 and influence estrogen receptor-regulated transcription (Zhang et al., 2011). Protein sequence analysis revealed a bipartite nuclear localization sequence near the N-terminal end and a number of serine and arginine (SR)-rich repeats (Figure S1A). Interestingly, proteins which contain arginine and serine dipeptide repeats, such as those in the RS and RS-related families, play important roles in constitutive and alternative splicing by participating in spliceosome formation and splice site selection (Kavanagh et al., 2005). While no formal RNA recognition motifs are present in ARGLU1, it can be classified as an ‘SR-like’ protein (Manley and Krainer, 2010). In the glutamate-rich domain we located two non-classical LXXLL motifs, LLXXL (172-176 aa) and LXXIL (201-205 aa) (Figure S1A). LXXL/IL motifs are canonical interaction sequences found in many NRs coactivators that facilitate binding to the AF2 domains of the receptors.

### ARGLU1 is constitutively nuclear and ubiquitously expressed

We next examined the intracellular distribution of ARGLU1 and compared it to that of GR in the presence and absence of a synthetic GC, dexamethasone (Dex), in HEK293 cells. Figure 1E shows ARGLU1 was constitutively nuclear with a distinct punctate pattern. As expected, GR was localized primarily in the cytoplasm but translocated into the nucleus and co-localized with ARGLU1 when Dex was added (Figure 1E).

To examine the tissue distribution of *Arglu1*, we analyzed its mRNA expression in 53 different C57Bl/6 mouse tissues by QPCR. *Arglu1* is ubiquitously expressed with the highest level of expression in the central nervous system. The mRNA expression of *Gr* and that of *Arglu1* were found to be highly correlated with only a few exceptions (i.e., uterus and pancreas (GR); testis (ARGLU1), Figure 2A).

**Figure 2:**
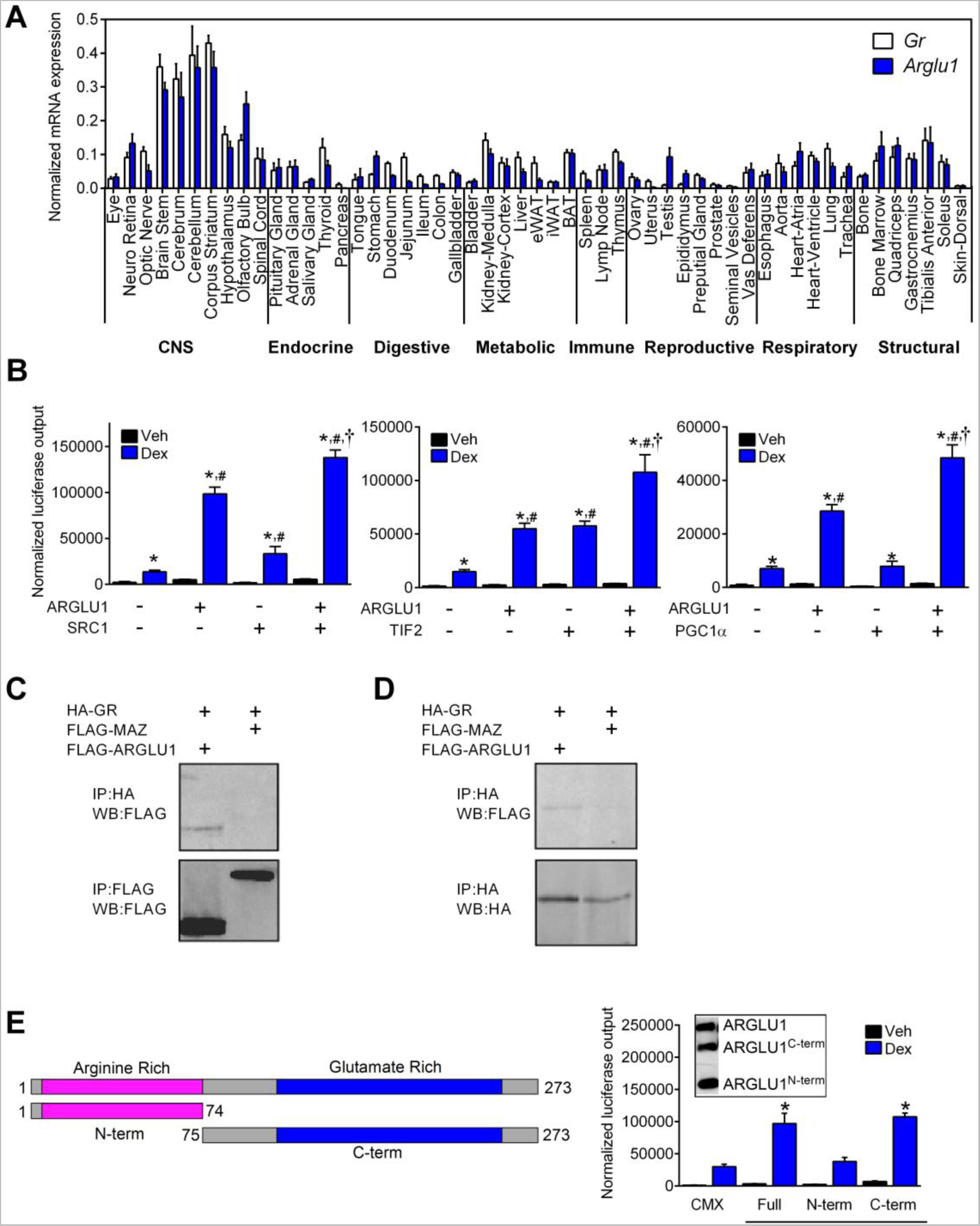
ARGLU1 is enriched in the CNS and coactivates GR via its C-terminal domain. (**A**) Tissues from male C57Bl/6 mice 4 months of age were collected. Ovary and uterus were from female mice. RNA was extracted and samples were DNase treated, reverse transcribed, and analyzed by QPCR (efficiency-corrected *Δ*Ct method). *36b4* was used as an endogenous reference RNA normalizer gene. Data represent the mean ± SD of individual QPCR well. Each tissue was pooled from at least 2 animals. eWAT, epididymal white adipose tissue; iWAT, inguinal white adipose tissue; BAT, brown adipose tissue. (**B**) HEK293 cells were transiently transfected with GAL4-GR, UAS-luc and ARGLU1 alone or in combination with known NR coactivators: SRC1, TIF2 and PGC1α; followed by administration of EtOH (Veh) or 100 nM Dex for 16 hrs. Data represent the mean ± SD (n = 3). *p < 0.05 Veh vs. Dex for respective condition; #p < 0.05 relative to empty vector (CMX)-Dex; † p < 0.05 vs ARGLU1-Dex; ANOVA followed by Newman-Keuls test. (**C-D**) HEK293 cells were transfected with HA-GR and either FLAG-ARGLU1 or FLAG-MAZ and treated with 100 nM Dex. Whole cell lysates were used to IP with anti-HA (**C**) or anti-FLAG (**D**) antibodies. Mouse anti-FLAG or rabbit anti-HA antibodies were used to confirm protein expression by Western blot. IP, immunoprecipitation; IB, immunoblot; MAZ, MYC-associated zinc finger protein, used as a negative control. (**E**) Schematic diagram of ARGLU1 truncations (left) and co-transfection assay of GAL4-GR/UAS-luciferase with 15 ng of ARGLU1 truncations in HEK293 cells treated with EtOH (Veh) or 100 nM Dex (right). CMX was used as a control. Data represent the mean ± SD (n = 3). Inset: FLAG-tagged full length ARGLU1 or the indicated truncation mutants were co-expressed in HEK293 cells. *p < 0.05 vs. CMX-Dex; ANOVA followed by Newman-Keuls test. See also Figure S3.

### ARGLU1 is a general NR coactivator and acts in concert with other coactivators

To test whether ARGLU1 can coactivate other members of the NR superfamily, we used the GAL4-NR-LBD/UAS-luciferase system. ARGLU1 increased the ligand-induced transcriptional activity of several steroid receptors (GR, ERα, MR, PR, ERβ) as well as a number of RXR heterodimer partners (PPARα,β,γ, LXRα,β and VDR), while no effect was observed for GAL4-FXR (Figure S2). Interestingly, ARGLU1 appeared to have the greatest effect on ligand-dependent coactivation of GAL4-GR and GAL4-ERα.

Nuclear receptor coactivators are usually present as part of multiprotein complexes and are known to potentiate each other’s activity (Lee et al., 2002). Here, we examined whether ARGLU1 can act together with other nuclear receptor coactivators to increase GR transcriptional activity. As expected, co-transfecting ARGLU1 with the GAL4-GR/UAS-luciferase system in HEK293 cells led to a ligand dependent increase in luciferase activity (Figure 2B). Similar results were observed when SRC1, TIF2 and PGC1α were added individually with the GAL4-GR/UAS-luciferase system (Figure 2B). When each of these coactivators was co-expressed with ARGLU1, the increase in ligand induced luciferase activity was additive (Figure 2B), implying that ARGLU1 and these other coregulatory proteins may form a complex with GR to regulate gene expression.

### ARGLU1 and GR co-immunoprecipitate from cells

To test our proposed model of ARGLU1 as a coactivator, we performed Co-IP assays using total cell extracts obtained from HEK293 cells transiently transfected with tagged versions of GR (HA), ARGLU1 (FLAG) or MAZ (a nuclear protein used as a negative control, FLAG). ARGLU1 was detected in the GR pulldown experiment, whereas, the negative control MAZ was not; yet, both proteins were abundantly expressed (Figure 2C). The reverse pull down with ARGLU1 confirmed that this interaction was specific to GR (Figure 2D).

### The C-terminal glutamate-rich domain of ARGLU1 is responsible for GR coactivation

With this protein interaction confirmed, we next determined the molecular domain of ARGLU1 responsible for GR coactivation using co-transfection assays with the GAL4-GR/UAS-luciferase system and truncation mutants of ARGLU1. These studies revealed that the construct containing only the C-terminus (ARGLU1^C-term^) retained the ability to coactivate GR, comparable to that of the full-length protein; whereas, the construct containing only the N-terminus (ARGLU1^N-term^) did not (Figure 2E). Equal protein expression of the truncation mutants was confirmed by Western blotting (Figure 2E, inset). To identify whether the LXXL/IL motifs we found in the C-terminal end (Figure S1A) were essential for GR coactivation, we mutated the leucine/isoleucine residues to alanine and tested these mutants in the co-transfection assay. We found no significant difference in GR coactivation when either NR-box was mutated suggesting that these motifs alone are not essential for ARGLU1 mediated transactivation of GR (Figure S3).

### ARGLU1 interacts with splicing factors and binds RNA at the arginine-rich N-terminal domain

To gain a more complete picture of the ARGLU1 protein interaction landscape, FLAG-ARGLU1 was purified from stable mammalian HEK293 cells and interacting proteins were identified by mass spectrometry (MS). ARGLU1 was found to interact with many proteins known to be implicated in RNA processing and splicing. Numerous members of the snRNP (small nuclear ribonucleoproteins), hnRNP (heterogeneous nuclear ribonucleoproteins), PRPF (pre-mRNA processing factors), RBM (RNA binding motif proteins), SFRS (splicing factor, arginine/serine-rich) and SF (splicing factors) protein families make up the ARGLU1 interactome (Figure 3A).

**Figure 3:**
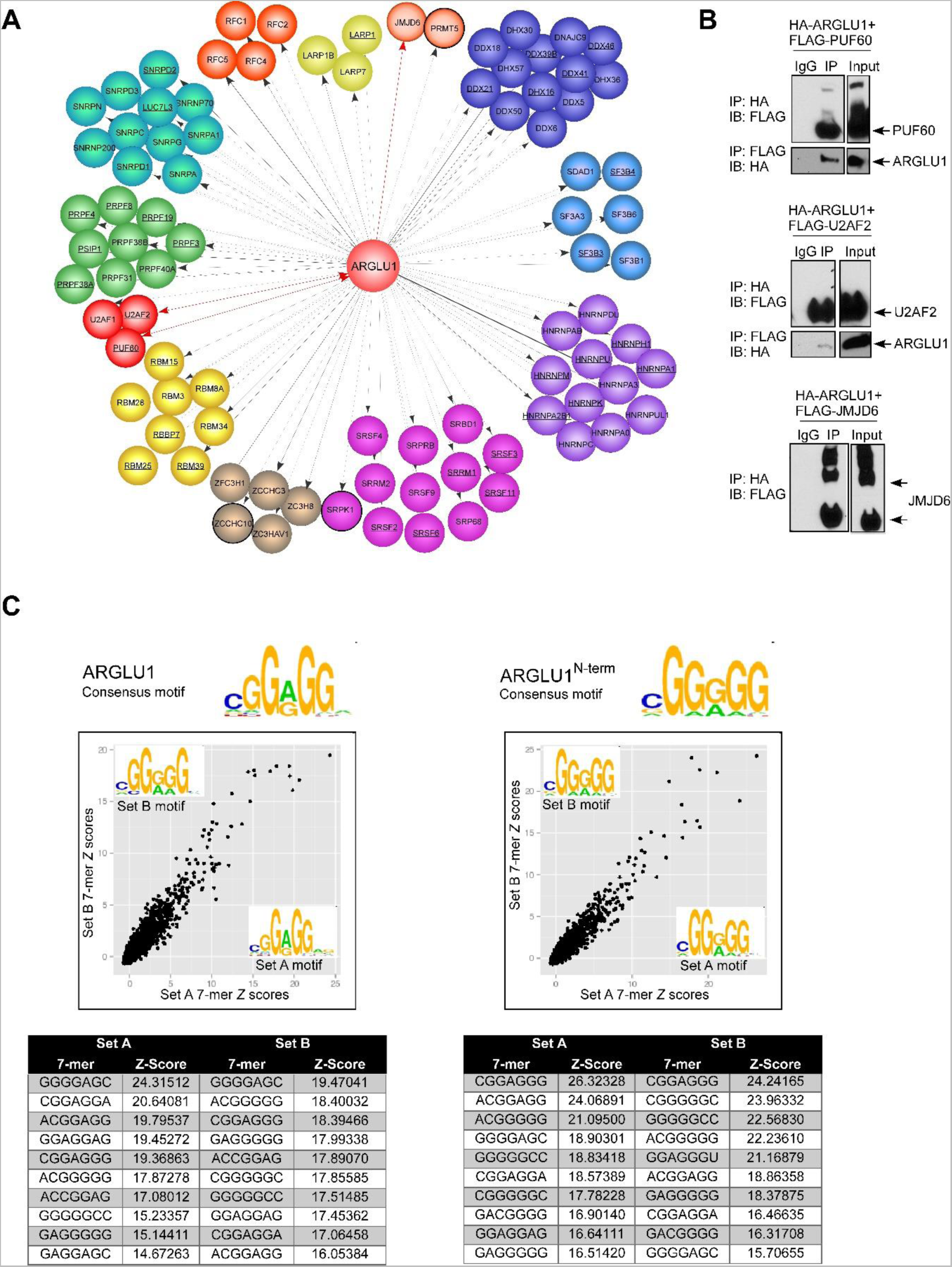
ARGLU1 interacts with splicing factors and binds to RNA. (**A**) ARGLU1 protein interaction network. HEK293 cells stably expressing ARGLU1 were incubated with FLAG M2 agarose and interacting proteins were tryptically digested and analyzed by MS. Cytoscape was used to generate the ARGLU1 protein interactome. Proteins which were pulled down in the BioID assay are underlined. Red lines depict interactions validated by co-IP experiments. Previously reported ARGLU1 interactors identified by GeneMANIA (Warde-Farley et al., 2010), have black borders. Pull-down experiments were repeated at least 3 times. (**B**) Co-IP of HA-ARGLU1 and selected factors (FLAG-PUF60, FLAG-U2AF2 or FLAG-JMJD6) identified by MS in (**A**) from whole cell lysates. FLAG-JMJD6 runs as a multimer on a gel. Reverse IPs with the FLAG antibody led to the HA-ARGLU1 being pulled down with FLAG-PUF60 andFLAG-U2AF2. IP, immunoprecipitation; IB, immunoblot. (**C**) RNAcompete results for GST-ARGLU1 and GST-ARGLU1^N-term^ (N-terminus intact). The scatter plots depict correlations between 7-mer Z-scores for set A and set B. Spots corresponding to enriched 7-mers are in the top right corner. Logos for consensus RNA binding motifs, averagedfrom set A and set B, were generated and shown at the top of the panel. The top ten 7-mers bound by the various ARGLU1 proteins (and corresponding Z-scores) are shown at the bottom of the panel.

To limit the possibility that transient interactions with ARGLU1 could be missed by standard IP-MS, we also performed proximity dependent biotin identification (BioID). Full length ARGLU1 was fused to a promiscuous *E. Coli* biotin protein ligase mutant (BirA R118G, BirA*), which upon introduction into *Flp***-**In™ *T-REx*™ 293 *cells* and addition of biotin results in biotinylation of nearby proteins (Kwon et al., 2002; Roux et al., 2012). Biotinylated proteins were purified by streptavidin pull down and identified by MS. BioID did not reveal additional novel interactors but confirmed our FLAG pull down results showing that ARGLU1 interacts with various proteins involved in constitutive and alternative splicing (Figure 3A).

The MS results were confirmed with Co-IP experiments in which ARGLU1 was found to interact with PUF60, U2AF2 and JMD6 (Figure 3B). PUF60 and U2AF2 (also known as U2AF65) are two factors which participate in intron-exon junction recognition by the spliceosome, whereas JMJD6 is an enzyme which has been found to hydroxylate U2AF2 and alter its activity (Hastings et al., 2007; Valcarcel et al., 1996; Webby et al., 2009).

Since ARGLU1 interacts with numerous splicing factors we wanted to determine if it can directly bind specific RNA sequences. To do this, we analyzed full-length GST-ARGLU1 and the two truncated proteins, GST-ARGLU1^N-term^ (N-terminus intact) and GST-ARGLU1^C-term^ (C-terminus intact) using RNAcompete (Ray et al., 2009; Ray et al., 2013). Purified GST-tagged proteins were incubated with a custom designed RNA pool comprised of ∼240,000 short RNA sequences, 30-41 nucleotides in length. RNAs bound to ARGLU1 were purified, labeled with either Cy3 or Cy5, and hybridized onto custom Agilent 244 K microarrays.

Computational analysis of the microarray data found full length ARGLU1 preferentially bound to CGG(A/G)GG type k-mers (Figure 3C). Interestingly, ARGLU1^N-term^ bound similar G-rich motifs (Figure 3C). In contrast, although RNA binding for ARGLU1^C-term^ was observed, no consistent and clear motif was identified (data not shown). Thus, ARGLU1 interacts with a specific RNA motif via its N-terminal domain.

### Regulation of basal and GC-mediated transcription in N2a cells by ARGLU1

In light of the high expression of *Arglu1* in the CNS (Figure 2A), we proceeded to characterize the role of ARGLU1 in mouse neuroblastoma Neuro-2a (N2a) cells. Given that over 90% of multiexonic genes expressed in the brain are alternatively spliced (Pan et al., 2008), and that we found ARGLU1 interacts with proteins of the AS machinery, we were curious to explore whether ARGLU1 has a dual role in GC-mediated transcription and AS in this cell system that has been previously used in this context (Calarco et al., 2009; Raj et al., 2011). To do so, RNA-seq was performed in N2a cells with *Arglu1* knockdown in the presence or absence of Dex. ARGLU1 knockdown resulted in a >90% reduction in mRNA and protein with no effect on *Gr* expression (Figure 4A,B).

**Figure 4:**
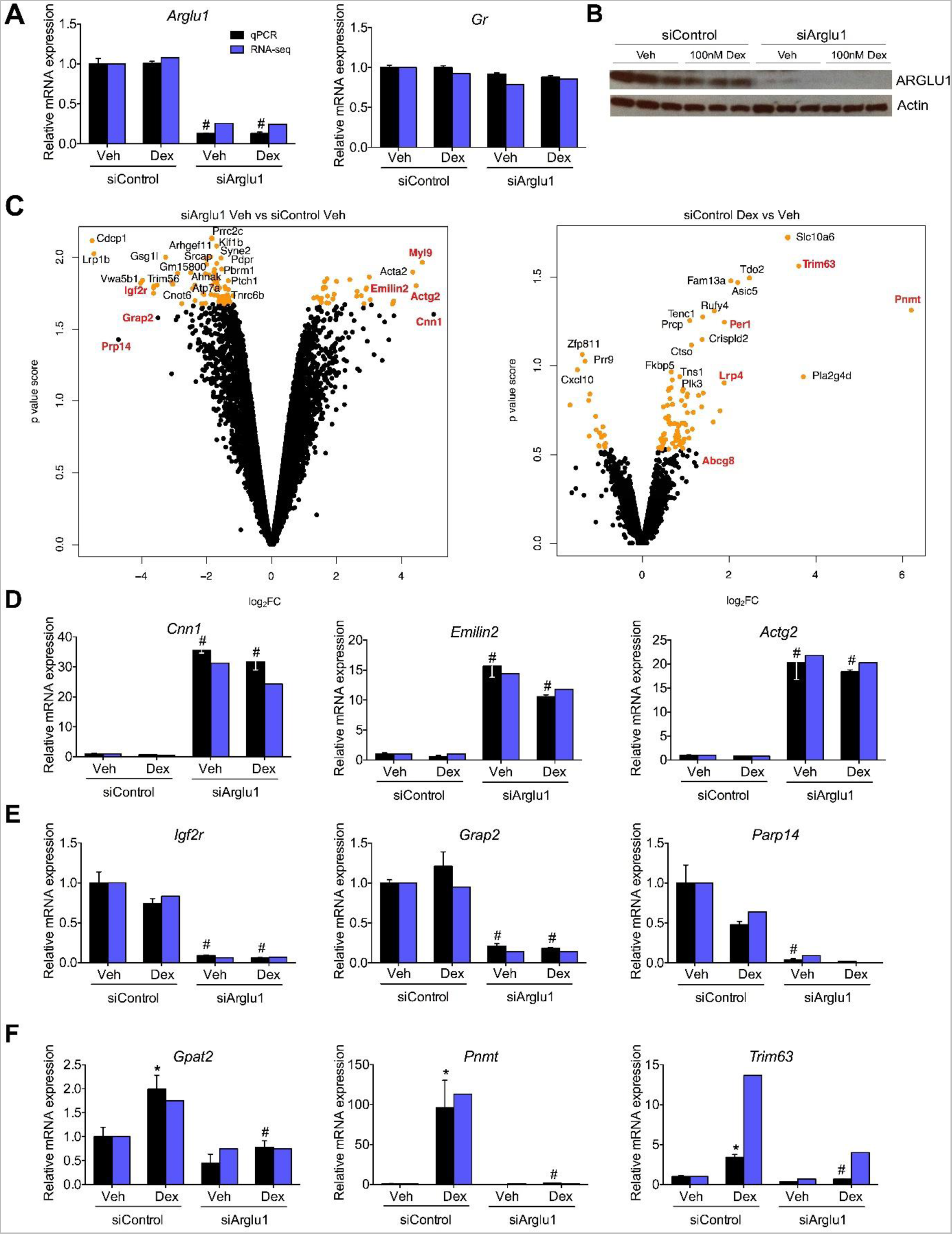
ARGLU1 influences basal and GC-induced gene expression in Neuro-2a cells. Neuro-2a cells were transfected with 30 pmol of siControl and siArglu1using RNAiMax for 48 hrs and then treated with vehicle (EtOH) or 100 nM Dex for 4 hrs before RNA or protein extraction. Quantitative PCR of (**A**) *Arglu1* and *Gr* mRNA normalized to cyclophilin. (**B**) ARGLU1 protein expression in whole cell lysates analyzed by Western blot (n = 3). (**C**) Volcano plot of log_2_ fold change (log_2_FC) versus p value score (defined in methods from edgeR analysis) for differentially expressed genes. Orange - top 100 genes. Red – genes validated by QPCR. (**D-F**) QPCR validation of RNA-seq data was performed on non-pooled samples. Fold changes obtained from RNA-seq (cRPKM) are plotted for comparison. Basal mRNA expression changes (siControl vs siArglu1) of genes upregulated (**D**) and downregulated (**E**) following ARGLU1 knockdown. (**F**) Ligand-dependent gene expression changes (Dex vs Veh). Data represent the mean ± SEM (n = 3). *p < 0.05 vs respective Veh, #p < 0.05 vs siControl; ANOVA followed by Newman-Keuls test. See also Figure S4.

Analysis found 607 genes (out of 12,061 total, p<0.05) were differentially expressed following ARGLU1 knockdown (siControl vs siArglu1) in the absence of a GC ligand (see Figure 4, S4A, and Table S1). A subset of these genes were validated by QPCR (Figures 4D-E, S4B) and showed a strong correlation with RNA-seq data (r=0.934, Figure S4C).

Following validation, we searched for pathways enriched in ARGLU1 transcriptionally regulated genes using gene set enrichment analysis (GSEA). Knockdown of ARGLU1 (in the absence of Dex) upregulated pathways involved in mitochondrial respiration, ribosome function and peptide hormone synthesis (Figure S4D). In contrast, pathways involved in chromatin and DNA modifications and circadian rhythm were significantly downregulated upon ARGLU1 knockdown (Figure S4D).

Next, we examined the gene expression changes in response to Dex in the presence or absence of ARGLU1. We identified 7 Dex-responsive differentially expressed genes using edgeR (p<0.05, Table S1, siControl) and validated a sub-set by QPCR (Figure 4F). The fold-change in the expression of 6 of the 7 differentially expressed genes (Dex/Veh) was decreased by the absence of ARGLU1 suggesting that, in neuronal cells, ARGLU1 is potentiating the majority of the GC-induced transcriptional changes. Interestingly, ARGLU1 knockdown completely abolished Dex-mediated induction of one of the top hits *Pnmt* (Figure 4F). *Pnmt* is involved in epinephrine synthesis and is known to be a direct target of GCs in the adrenal gland (Evinger et al., 1992). Another direct target of GR, *Trim63*, an E3 ubiquitin ligase, also showed a significant decrease in Dex-mediated induction upon ARGLU1 knockdown (Azuma et al., 2010).

### Regulation of basal and GC-mediated alternative splicing in N2a cells by ARGLU1

Analysis of RNA-seq data showed ARGLU1 was essential for proper basal and GC-induced AS in N2a cells. Alternative splicing was assessed using an established bioinformatics pipeline (Irimia et al., 2014) from which the “percent spliced in” (PSI) metric was generated. For our analysis, only splicing events that showed changes in PSI (delta PSI, dPSI) of ≥15 were included. Knockdown of ARGLU1 in vehicle treated cells resulted in 1129 differential AS events (within 928 genes) compared to siControl (calculated as siArglu1 Veh PSI – siControl Veh PSI) (Table S2), indicating that ARGLU1 is necessary for establishing AS patterns in N2a cells. The majority of AS events affected by ARGLU1 belonged to the simple and complex cassette exon category (Figure 5A). When comparing the direction of the change in cassette exons within the differential AS events, loss of ARGLU1 promoted exon inclusion or skipping in 44% and 56% of events, respectively (Figure 5A). One-step RT-PCR was used to validate 20 AS events showing differential inclusion levels upon ARGLU1 knockdown (Figures 5C, S5A). The difference in the PSI values calculated from RNA-seq and one-step RT-PCR were highly correlated (r=0.947, Figure S5B). Similar results were observed with two independent siRNAs targeting ARGLU1 (Figure S6).

**Figure 5:**
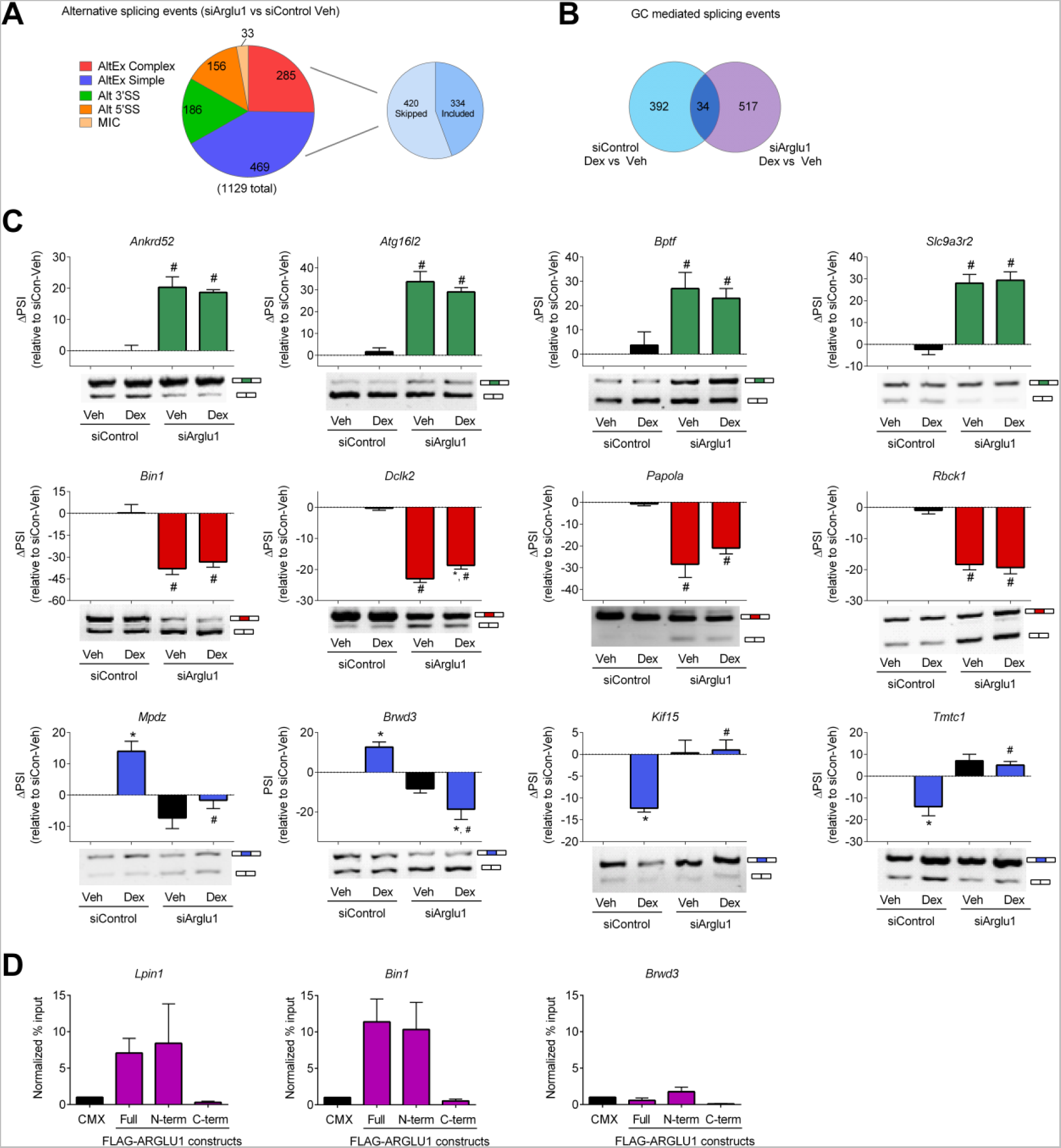
ARGLU1 influences basal and GC-induced alternative splicing in Neuro-2a cells. (**A**) Classification of ARGLU1-regulated events. Pie charts showing the distribution of alternative spliced events following ARGLU1 knockdown in N2a cells (in the absence of ligand stimulation). AltEx refers to simple and complex cassette exon events; 3′SS, alternative 3′ splice site; 5′SS, alternative 5′ splice site; MIC, microexon. Pie chart inset: distribution of skipped or included exons within the combined simple and complex cassette exon category with ARGLU1 knockdown. (**B**) Venn diagram showing overlap in alternative splicing events (PSI ≥ 15) by Dex in the presence (siControl) or absence of ARGLU1 (siArglu1). (**C**) Splicing events with a PSI ≥15 were validated using one-step RT-PCR. Representative image is shown below the Image J quantification of the blot. Data represent the mean ± SEM (n=3). *p<0.05 vs respective Veh, #p<0.05 vs respective siControl; ANOVA followed by Neuman-Keuls test. (**D**) RNA immunoprecipitation in N2a cells with full length FLAG-tagged ARGLU1 and the indicated truncation mutants was performed on genes containing one or more putative ARGLU1 binding sites identified by visual examination within ± 300 bp of the alternatively spliced exon or a negative control gene that undergoes splicing but is not dependent on ARGLU1. RNA was purified and quantified using One-Step QPCR (Froggabio) with primers spanning exon-intron junctions to look at pre-mRNA binding by ARGLU1. Signals were corrected for input and expressed as normalized fold enrichment over CMX-transfected cells. See also Figure S5 and S6.

Next, we looked for Dex-responsive AS events. There were a total of 426 AS events that showed absolute dPSI of ≥15 in response to Dex in the siControl group (calculated as siControl Dex PSI – siControl Veh PSI) (Figure 5B; Table S2). Remarkably, 392/426 (92%) of these events were dependent on ARGLU1. These data thus show that ARGLU1 is essential for maintaining proper GC-induced AS in N2a cells

To examine whether ARGLU1 can bind *in vivo* to the pre-mRNA of alternatively spliced genes, we manually identified potential ARGLU1 binding sites ±300 bp surrounding the alternatively spliced exons by searching the pre-mRNA sequences of AS genes for GGAGG or GGGGG. Using RNA immunoprecipitation, we observed binding of full length ARGLU1 and ARGLU1^N-term^ to the pre-mRNA of *Lpin1* and *Bin1* (Figure 5D). ARGLU1^C-term^ did not show any binding above background levels (Figure 5D). Neither full length ARGLU1 nor ARGLU1^N-term^ showed any binding to the pre-mRNA of *Brwd3*, a gene that does not contain a G-rich ARGLU1 binding motif but does undergo alternative splicing (Figure 5D).

### ARGLU1 regulates gene expression and AS on distinct genes within similar networks

It is widely accepted that splicing and transcription are temporally coupled (Naftelberg et al., 2015). To explore whether ARGLU1 facilitates the coupling of these two processes, we compared the genes affected by ARGLU1 in each category. Interestingly, out of the 607 transcriptionally regulated and 928 alternatively spliced genes that were affected by ARGLU1 knockdown (siArglu1 vs siControl), only 71 genes were overlapping (Figure 6A, Table S3).

**Figure 6:**
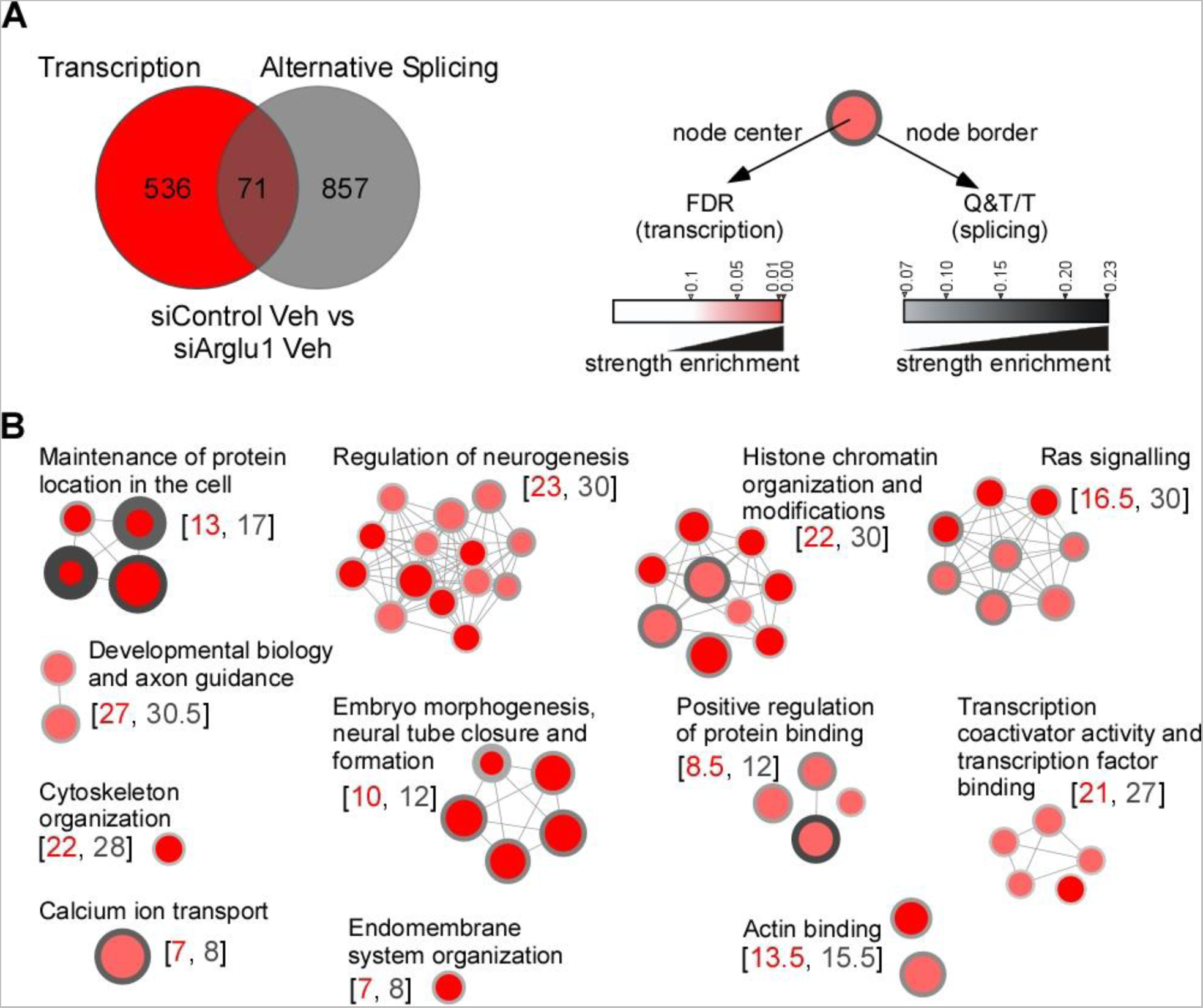
ARGLU1 regulates different genes by transcription and AS, but these genes belong to overlapping functional pathways. (**A**) Venn diagram showing limited overlap in genes alternatively spliced (PSI ≥ 15) and transcriptionally regulated (siArglu1/siControl log2 ratio ≤-0.81 or ≥0.81) by ARGLU1 in N2a cells in the absence of a ligand. Genes having more than one splicing event were only counted once. (**B**) Pathways demonstrating overlap between those basally regulated by ARGLU1 in alternative splicing vs. transcription (siArglu1 vs siControl). Node size is proportional to the number of differentially expressed genes in a pathway (corrected by the size of the pathway). All these pathways have a significant enrichment FDR equal or less than 0.05 for both the alternative splicing and transcription gene lists. Related pathways are grouped together under a common label to form modules. The numbers in brackets correspond to the median number of genes for transcription (left) and alternative splicing (right) contained in each pathway module. FDR, false discovery rate. Q&T/T, number of genes in the overlap for each pathway corrected by the size of the pathway.

Strikingly, although differentially expressed and alternatively spliced genes were largely unique, they were frequently enriched within the same pathways including ‘regulation of neurogenesis’ and ‘histone chromatin organization and modifications’ (Figure 6B). These data suggest that although ARGLU1 basally regulates the AS and expression of distinct genes, these genes share a similar functional classification.

### Loss of ARGLU1 is embryonic lethal in mice

To investigate the physiologic role of ARGLU1 *in vivo*, we attempted to generate a whole-body ARGLU1 knockout mouse. Heterozygous Arglu1 mice (*Arglu1^+/-^*also known as *Arglu1^tm1a/+^*) were generated by morula aggregation using commercially available ES cell lines (EUCOMM, details provided in the Extended methods and Figure S7). Male and female heterozygous *Arglu1^+/-^* mice appeared phenotypically normal.

Breeding of heterozygous *Arglu1^+/-^* mice did not produce any homozygous knockout mice (out of 200 pups) suggesting that *Arglu1^-/-^* mice are embryonic lethal. To begin to identify the developmental defects, we performed timed pregnancy experiments. Genotyping of the yolk sacs detected *Arglu1^-/-^* embryos at E9.0 and E9.5 but not at E12.5. Gross morphologic examination of these mutant embryos revealed an overall developmental delay of ∼0.5 days at E9.0 and E9.5 (Figure S7E). This mid-gestation lethality is consistent with a generalized growth defect and demonstrates that ARGLU1 is indispensable for embryonic development before E9.5.

### ARGLU1 is important for proper CNS development in zebrafish

The high evolutionary conservation of ARGLU1 allowed us to utilize the zebrafish *Danio rerio* as a second model organism to study ARGLU1’s role *in vivo*. Interestingly, unlike mammalian ARGLU1, which is encoded by only one gene, zebrafish possess two ARGLU1 paralog genes encoded on separate chromosomes. The presence of two paralogs in zebrafish is consistent with the teleost specific whole genome duplication that occurred during vertebrate evolution (Howe et al., 2013). The sequence identities of ARGLU1a and ARGLU1b to human ARGLU1 are 84% and 87%, respectively. The expression of *arglu1a* and *arglu1b* was examined by whole-mount RNA *in situ* hybridization at 24 and 48 hours post-fertilization (hpf). Diffuse expression of both isoforms was observed throughout the developing brain at each time point (Figure 7A-D). These results are consistent with the high CNS expression of *Arglu1* in mice (Figure 2A).

**Figure 7:**
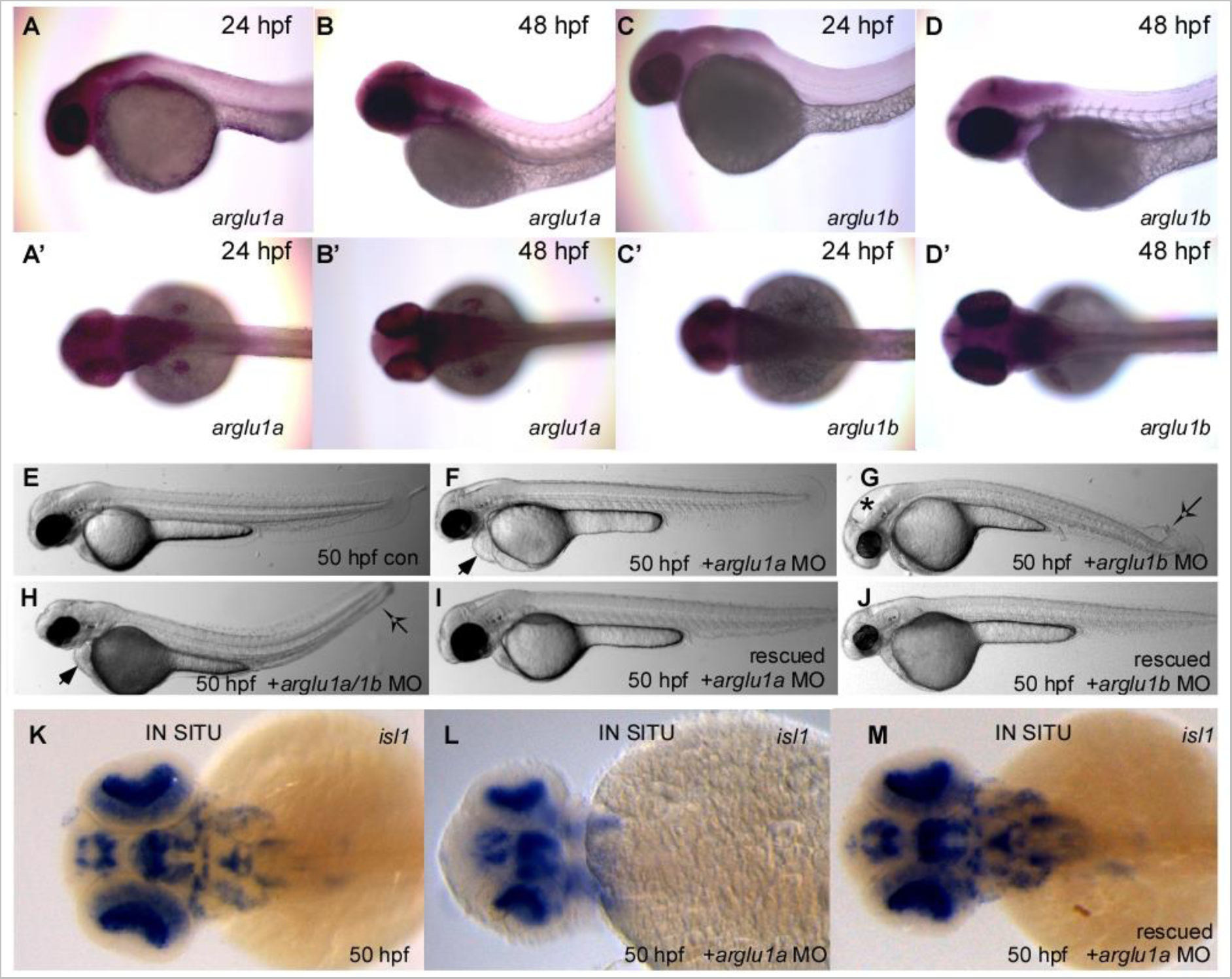
ARGLU1 regulates vertebrate nervous system development. (**A–D**) RNA *in situ* hybridization assays monitoring *arglu1a* and *arglu1b* expression in the developing zebrafish embryo. Expression of *arglu1a* and *arglu1b* at 24 hpf (**A, C** and **A’, C’**) and 48 hpf (**B, D** and **B’, D’**), lateral and dorsal views, respectively. (**E**) p53 morpholino antisense oligonucleotide (MO) control-injected embryo 50 hpf. (**F**) p53 + arglu1a MO-injected embryo 50 hpf. (**G**) p53 + arglu1b MO-injected embryo50 hpf. (**H**) p53 + arglu1a MO + arglu1b MO-injected embryo at 50 hpf. Symbols represent heart edema (), expanded brain ventricle (*) and curved body axis). (**I**) p53 + arglu1a MO injected embryo, rescued through coinjection of *arglu1a* mRNA. (**J**) p53 + arglu1b MO injected embryo, rescued through coinjection of *arglu1b* mRNA. (**K-M**) RNA *in situ* hybridization assays monitoring islet-1 expression in 50 hpf embryos. p53-MO-injected control (**K**), arglu1a morphant embryo (**L**), and arglu1a morphant embryo rescued through coinjection of *arglu1a* mRNA (**M**) are shown. Hpf, hours post fertilization. See also Figure S7.

To examine the effect of loss of ARGLU1 proteins in zebrafish, we injected single cell-staged embryos with translation blocking morpholinos (MO) targeting *arglu1a* and *arglu1b*, alone or in combination. To minimize the off-target effects of morpholino injections mediated by the p53 apoptotic pathway, p53 MO was co-injected (Robu et al., 2007). At 50 hpf, p53 MO injected control embryos appeared phenotypically normal (Figure 7E); whereas, more than 70% of arglu1a MO-injected embryos exhibited heart edema and decreased brain size (n=85, Figure 7F). Interestingly, arglu1b MO-injected animals (more than 90%) showed a different morphant phenotype, with expanded brain ventricles and a curved body axis (n=85, Figure 7G). Co-injection of arglu1a and 1b MO together led to a more severe phenotype, where animals exhibited abnormal brain development, heart defects and a curved body axis (n=30, Figure 7H). All embryos which had the morphant phenotype showed impaired movement without lethality up until the feeding stage (8-9 days) where they were unable to feed and died. To confirm that the observed results were not due to off-target effects of MO injection, we performed rescue experiments by co-injecting the arglu1a and arglu1b MOs with *in vitro* transcribed *arglu1a* or *arglu1b* mRNA, respectively (Figure 7I-J). *arglu1a* and *arglu1b* mRNA was able to rescue the morphant phenotypes in 65% of the embryos (n=60, Figure 7I-J).

To better delineate the brain defect in arglu1a MO embryos, we performed RNA *in situ* hybridization for *islet-1*, a LIM homeodomain-containing transcription factor which acts as an early marker of neuronal differentiation. At 50 hpf, p53 MO injected control animals showed the characteristic *islet-1* expression pattern in the brain (Figure 7K), whereas arglu1a MO injected embryos had strikingly reduced *islet-1* expression, indicating possible disruption of neuronal differentiation (Figure 7L). Validating the on-target activity of the MO, co-injection of *arglu1a* mRNA rescued expression of *islet-1* in these embryos (Figure 7M). Overall, our results suggest that ARGLU1 proteins in zebrafish play a critical role in proper brain development.

## Discussion

### ARGLU1 is an evolutionarily conserved NR coactivator and AS modulator

We detail the discovery and characterization of ARGLU1 as a highly conserved dual function protein working as both a GR coactivator and AS effector. ARGLU1 represents a new member within the diverse family of NR coregulatory proteins, since it shares no sequence similarity among other coregulators identified to date and, despite its ability to bind RNA, contains no known mammalian RNA binding domains. Deletion of ARGLU1 was embryonic lethal in mice between E9.5 and E12.5, and resulted in severe developmental defects in the CNS and heart of zebrafish. Biochemical studies and unbiased proteomics analyses determined that ARGLU1 interacts with GR through its C-terminal domain; whereas, the N-terminal domain mediates interactions with splicing factors and contributes to both basal and GC-dependent splicing outcomes. Remarkably, only 7.5% of the genes differentially alternatively spliced by ARGLU1, were also transcriptionally regulated by ARGLU1. These findings strongly implicate ARGLU1 as a dual-function protein, utilizing its two distinct structural domains to regulate gene expression at two levels (transcription and pre-mRNA splicing) to increase protein diversity within selected regulated pathways basally and in response to hormonal ligands.

Consistent with many NR coregulators, ARGLU1 coactivates multiple receptors in both a ligand dependent and independent fashion. In agreement with our discovery of ARGLU1 as a GR coactivator, Zhang and colleagues showed that ARGLU1 interacts with a mediator complex subunit and is required for amplifying ERα–mediated gene transcription. Among the receptors that showed ligand dependence for ARGLU1, GR was the most highly dependent followed by ERα and several others to a much lower extent. The additivity in the transcriptional output ARGLU1 with other GR coactivators (SRC1, TIF2 and PGC1α) is consistent with its function as a coactivator. Such effects have been observed previously between coactivators and NRs (Lee et al., 2002; MacDonald et al., 2001) and suggest that ARGLU1 and/or TIF2, SRC1 and PGC1α may form a multi-protein complex with GR to regulate gene expression.

The two domains of ARGLU1 have opposing charges with positively charged arginine residues at the N-terminus and an abundance of negatively charged glutamate residues at the C-terminus. Proteomic and RNAcompete analyses indicated that the N-terminal end was responsible for interacting with splicing factors and binding RNA. Our findings are also supported by a proteomics study of the human spliceosome that identified ARGLU1 (FLJ10154) as one of the proteins repeatedly co-purified with the U2 related spliceosomal complex (Hegele et al., 2012). Intriguingly, although ARGLU1 contains no consensus mammalian RNA binding motifs, the highly enriched segment of arginines at the N-terminus is consistent with the mechanism by which some viral proteins bind RNA (i.e., HIV Rev and Tat proteins) (Calnan et al., 1991). The CGG(A/G)GG rich motifs that were found to preferentially bind ARGLU1 by RNAcompete are similar in sequence to that preferentially bound by SRSF2 (Ray et al., 2013). Indeed, there is recent evidence that SRSF2 may play an opposing role to that of ARGLU1 in splicing (B.J. Blencowe, personal communication). From computational studies, it has been suggested that this GGAGG rich motif may also function as an intronic splicing enhancer that acts in a combinatorial manner to compensate for weak PY tracts (Murray et al., 2008). While binding to an intronic splicing enhancer would be expected to promote exon inclusion; in neuronal cells, ARGLU1 showed almost equal preference in promoting exon skipping and inclusion. The ability to promote both inclusion and exclusion is in line with recent evidence demonstrating that binding site location is only one factor contributing to splicing outcomes, and the interaction with other splicing factors in a gene-specific context is also key (Huelga et al., 2012; Han et al., 2011; Pandit et al., 2013).

### Hormone dependent AS occurs independent of transcriptional activation on a genome-wide scale

NR co-regulator studies have traditionally examined the interplay of NRs and their coactivators in the initiation of gene transcription and their interaction with chromatin; with much less attention paid to the post-transcriptional role of NR coregulators in signaling (Lonard and O'Malley, 2005). AS is a regulated process by which pre-mRNAs from a single gene are differentially spliced into several mRNAs. Splicing is orchestrated by a multi-protein complex composed of 5 small ribonucleoprotein particles and up to ∼300 splicing factors (Black, 2003; Martinez-Contreras et al., 2007). It is estimated that ∼90% of human pre-mRNAs are alternatively spliced (Pan et al., 2008; Wang et al., 2008). The regulation of AS is known to occur co-transcriptionally and/or post-transcriptionally depending on the gene (Cramer et al., 1997; Luco et al., 2011; Vargas et al., 2011).

In the context of NR signaling, steroid NRs and their coactivators have been implicated in AS with the idea that activated steroid hormone receptors control gene transcription and affect splicing decisions in a promoter-dependent manner by recruiting a set of transcriptional coregulators that participate in the splicing decisions of the newly formed transcripts (Auboeuf et al., 2007; Auboeuf et al., 2004; Auboeuf et al., 2002). For example, PPARγ coactivator, PGC1α, was shown to influence the AS of the fibronectin minigene when PPAR/RXR binding sites were present in the promoter (Monsalve et al., 2000). Similarly, coregulators of steroid receptors, CAPER and CoAA, were found to play a role in the pre-mRNA processing of various minigenes in response to ligand (Auboeuf et al., 2004; Dowhan et al., 2005; Iwasaki et al., 2001). In cells, CAPERα was found to coactivate the progesterone receptor and alter the splicing of endogenous *Vegf* (Dowhan et al., 2005). In these models, the influence of the coregulatory protein on splicing was intimately tied to transcriptional activation, generally in response to hormone through the incorporation of the MMTV promoter upstream of the minigene construct. These results invoke the kinetic model of co-transcriptional splicing in which activation of transcription by the NR is a co-requisite to its role in splicing.

In contrast, our data suggest that the above model is not valid for the endogenous regulation of genes by glucocorticoids and may be only valid in the context of mini-gene reporters. When examined on a genome-wide scale, we find that among the genes that were altered in response to Dex, there was no overlap between genes transcriptionally regulated and those that were alternatively spliced. To our knowledge, our study is the first to examine the global effect of a NR coregulatory protein on AS in response to GC signaling. Although these findings were unexpected based on historical minigene data, our observations are in agreement with a study in which an exon array was used to probe the role of ERα and ERβ in estradiol-induced AS (Bhat-Nakshatri et al., 2013). This study found that ∼67% of genes that were alternatively spliced after a 3 hr treatment were not transcriptionally altered (Bhat-Nakshatri et al., 2013). More recently, RNA-seq analysis of breast cancer cells treated with a synthetic progestin found 254 genes alternatively spliced after a 6 hr treatment, 74% of which were not transcriptionally regulated (Iannone et al., 2015).

Overall, these data support two distinct functional roles for ARGLU1, one involved in transcription and the other in AS. It is not yet clear whether the developmental defects observed in mice and zebrafish with loss of ARGLU1 are due to the loss of its transcriptional or splicing role, but we anticipate that both will be important based on our bioinformatics analysis in neural cells that found developmental pathways were enriched in both the transcriptionally regulated and alternatively spliced gene lists. The degree to which ARGLU1 is interfacing specifically with GR to regulate genes is, as yet, not clear. However, we do note that of the genes significantly induced by Dex in N2a cells (siControl), the magnitude of induction was diminished for 6 out of 7 genes by the absence of ARGLU1. Likewise, of the 426 splicing events that were alternatively spliced in response to Dex, 392 (92%) were dependent on ARGLU1. Together, these data suggest that in N2a cells, ARGLU1 strongly influences GC signaling outcomes and remarkably, changes were largely at the level of alternative splicing. These data suggest that stress hormone-induced AS is a further layer of gene regulation that is independent of transcriptional activation. Because the brain is prone to large changes in alternative splicing, this finding may be particularly important with respect to stress-induced cognitive dysfunction and provide a new angle from which to explore the molecular mechanisms of hormone function in disease states.

## Experimental Procedures

For procedures listed below, additional details are available in the Supplemental Experimental Procedures.

### High-throughput expression screen and transfection assays

Electromax^TM^ DH10B^TM^ E.Coli cells (Invitrogen, Carlsbad, CA) were transformed with 10 ng of normalized human brain cDNA library, diluted and grown overnight in deep 96-well plates (Sigma, St. Louis, MO). The following morning, plasmid DNA was isolated from the bacterial cultures using the GenElute^TM^ HP 96 well plasmid midiprep kit (Sigma). Screening was performed by transfecting pools of the extracted plasmid DNA into HEK293 cells using calcium phosphate in the presence of indicated controls (Makishima et al., 1999). Six hours post-transfection, cells were treated with vehicle (ethanol) or 300 nM cortisol. Cells were harvested 14–16 hrs later for luciferase and β-galactosidase activity, as previously described (Makishima et al., 1999). Neuro-2a (N2a) cells were transfected in suspension with siRNA against *Arglu1* (D-057082-02; 5’-GCCAAACGCAUCAUGGAAA-3’) or with the non-targeting Control siRNA (siGENOME Non-Targeting siRNA Pool #2; D-001206-14-05) using RNAiMax as per manufacturer’s instructions.

### Confocal microscopy

HEK293 cells were co-transfected with Cherry-hARGLU1 and GFP-hGR. After 48 hours, cells were treated with either vehicle (ethanol) or 100 nM Dex for 4 hours and then fixed using 4% PFA (pH 7.4). Vectashield (Vector laboratories, Burlingame, CA) mounting media with DAPI was used to mount the coverslips on glass slides and visualized with a LSM 700 Zeiss confocal laser scanning microscope (Jena, Germany).

### Animal experiments

Three targeted ES cell lines heterozygous for the knockout-first-reporter tagged *Arglu1^tm1a(EUCOMM)Hmgu^* allele were obtained from European Conditional Mouse Mutagenesis Program (EUCOMM). The generation and breeding of the transgenic animals is described in the extended experimental procedures. C57Bl/6 mice used for tissue collection were housed in a temperature and light-controlled environment and fed *ad libitum* the 2016S diet (Harlan Teklad Mississauga, ON, Canada). Zebrafish were maintained at 28.5°C on a 14/10 hour light/dark cycle and staged relative to hours post-fertilization. All animal studies were performed according to the recommendations of the Faculty of Medicine and Pharmacy Animal Care Committee at the University of Toronto (Toronto, ON).

### Zebrafish morpholino injections and *in situ* hybridization

Translation-blocking morpholinos (MO) targeting zebrafish *arglu1a*, *arglu1b* and *p53* genes were purchased from Gene Tools, LLC (Philomath, OR). Morpholinos were injected alone or in combination with the indicated rescue RNA into one-cell stage embryos using standard techniques. Whole mount *in situ* hybridization was carried out as previously described with the indicated antisense RNA probes (Hauptmann and Gerster, 1994). Rescue RNA and antisense RNA probes were synthesized using mMESSAGE mMACHINE (Ambion; Austin, TX). Embryos were imaged at indicated times after fertilization.

### RNA isolation, cDNA synthesis, and real-time QPCR analysis

Total RNA was extracted with RNA STAT-60 (Tel-Test Inc, Friendswood, TX), treated with DNase I (Roche, Laval, QC, Canada) and reverse transcribed into cDNA with random hexamer primers using the High Capacity Reverse Transcription System (ABI, Burlington, ON, CA). Real-time quantitative PCR reactions were performed on an ABI 7900 machine in a 384-well plate format. Relative mRNA levels were calculated using the comparative Ct method (Bookout et al., 2006). For the tissue library mRNA analysis, the efficiency-corrected ΔCt method was used.

### FLAG affinity purification and BioID assays

Protein-interaction complexes from HEK293 cells stably expressing FLAG-hARGLU1 were extracted and subjected to mass spectrometry analysis as previously described (Zhang et al., 2014).

### Co-immunoprecipitation assays and Western Blot analysis

HEK293 cells were grown in 100 mm plates and transiently transfected with 5 µg of the expression plasmid of interest. Forty eight hours later, cells were harvested and protein was extracted. Protein lysates were pre-cleared with protein G beads, split into 3 tubes and incubated with 5 µg of indicated antibodies. The next day, 50 μL of protein G agarose was added to the lysates and incubated on a rotator for 3-4 hrs at 4°C. Proteins were eluted in lithium dodecyl sulfate buffer (LDS, Invitrogen) and resolved on a 4-20% gradient SDS gel (Bio-Rad, Hercules, CA).

### RNAcompete studies

The procedure was performed as previously described (Ray et al., 2013). Briefly, GST-tagged ARGLU1 (20 pmol) and RNA pool (1.5 nmol) were incubated in 1 mL of binding buffer (20 mM HEPES pH 7.8, 80 mM KCl, 20 mM NaCl, 10% glycerol, 2 mM DTT, 0.1 mg/mL BSA) containing 20 mL of glutathione sepharose 4B beads (GE Healthcare, pre-washed 3 times in binding buffer) for 30 minutes at 4°C. Beads were then washed and the recovered RNA was hybridized to the Agilent 244K microarray used to generate the RNA pool. One-sided Z-scores were calculated for the motifs as previously described (Ray et al., 2013).

### RNA-seq studies and analysis

N2a cells were transfected with siGENOME siRNA (Dharmacon) using RNAiMAX reagent (Invitrogen). mRNA was pooled by treatment group (n=3/group) and mRNA enriched Illumina TruSeq V2 RNA libraries were prepared. Samples were sequenced at the Donnelly Sequencing Centre (University of Toronto) on Illumina HiSeq2500.

Transcriptome-wide alternative splicing and gene expression profiling were performed using a previously described workflow (Irimia et al., 2014).

### Statistical analysis

Data are presented as mean ± SD unless otherwise indicated. One-way ANOVA followed by the Newman-Keuls test was used to compare more than two groups. For comparison between two groups the unpaired Student’s t-test was performed. All tests were performed using GraphPad Prism6. *P*<0.05 was considered significant.

## Author Contributions

Conceptualization, L.M. and C.L.C.; Formal Analysis, V.V., S.G., and M.I.; Investigation, L.M., J.T., E.Z., M.R., S.G., M.I., D.R., R.P. and J.P.; Resources, S.A.; Writing – Original Draft, L.M. and C.L.C.; Writing – Review & Editing, L.M., V.V., M.I., D.R., R.P., G.B., H.K., B.J.B., S.A., and C.L.C.; Visualization, L.M., J.T., and V.V.; Supervision, G.B., T.R.H., H.K., B.J.B., S.A., and C.L.C.; Funding Acquisition, C.L.C.

## Acknowledgments

The authors wish to acknowledge the contribution of Matthew Chow (University of Toronto) for preliminary bioinformatics analyses, Dr. C.C. Hui (University of Toronto) for help in acquiring embryo images, Dr. Keith Pardee (University of Toronto) for critically reviewing the manuscript, and Evan Easton (University of Pittsburgh) for assistance with Southern blotting. Lilia Magomedova was funded by an Ontario Graduate Scholarship. We gratefully acknowledge funding from the Connaught New Investigator Award (University of Toronto, C.L.C.), the J.P. Bickell Foundation (C.L.C.), and the Natural Sciences and Engineering Research Council of Canada (RGPIN 03666-14 to C.L.C. and CGS-D to L.M.).

## Supplemental Tables (uploaded as excel files)

Table S1. N2a RNA-seq gene expression data set. N2a RNA-seq data for genes which showed significant differential gene expression following ARGLU1 knockdown (p<0.05 by edgeR analysis).

Table S2. N2a RNA-seq alternative splicing data set. Only events with dPSI of ≥ 15 for the indicated condition are shown. Event complexity refers to the percentage of reads that do not come from the C1A, AC2, C1C2 exon junctions. S is a simple event with ≤5% of total reads being complex, while C1, C2, and C3 have an increased proportion of complex reads. Other codes refer to additional types of AS events, where MIC refers to microexons (≤ 15 nucleotides).

Table S3. N2a genes showing transcription and splicing overlap following ARGLU1 knockdown. N2a RNA-seq data for genes which showed significant (p<0.05 by edgeR analysis) changes in basal expression level upon ARGLU1 knockdown and overlapping genes which showed splicing changes of PSI of ≥ 15 (calculated as siArglu1 PSI – siControl PSI). Event complexity refers to the percentage of reads that do not come from the C1A, AC2, C1C2 exon junctions. S is a simple event with ≤5% of total reads being complex, while C1, C2, and C3 have an increased proportion of complex reads. Other codes refer to additional types of AS events, where MIC refers to microexons (≤ 15 nucleotides). A total of 71 overlapping genes were identified.

## Supplemental Figures

**Figure S1:**
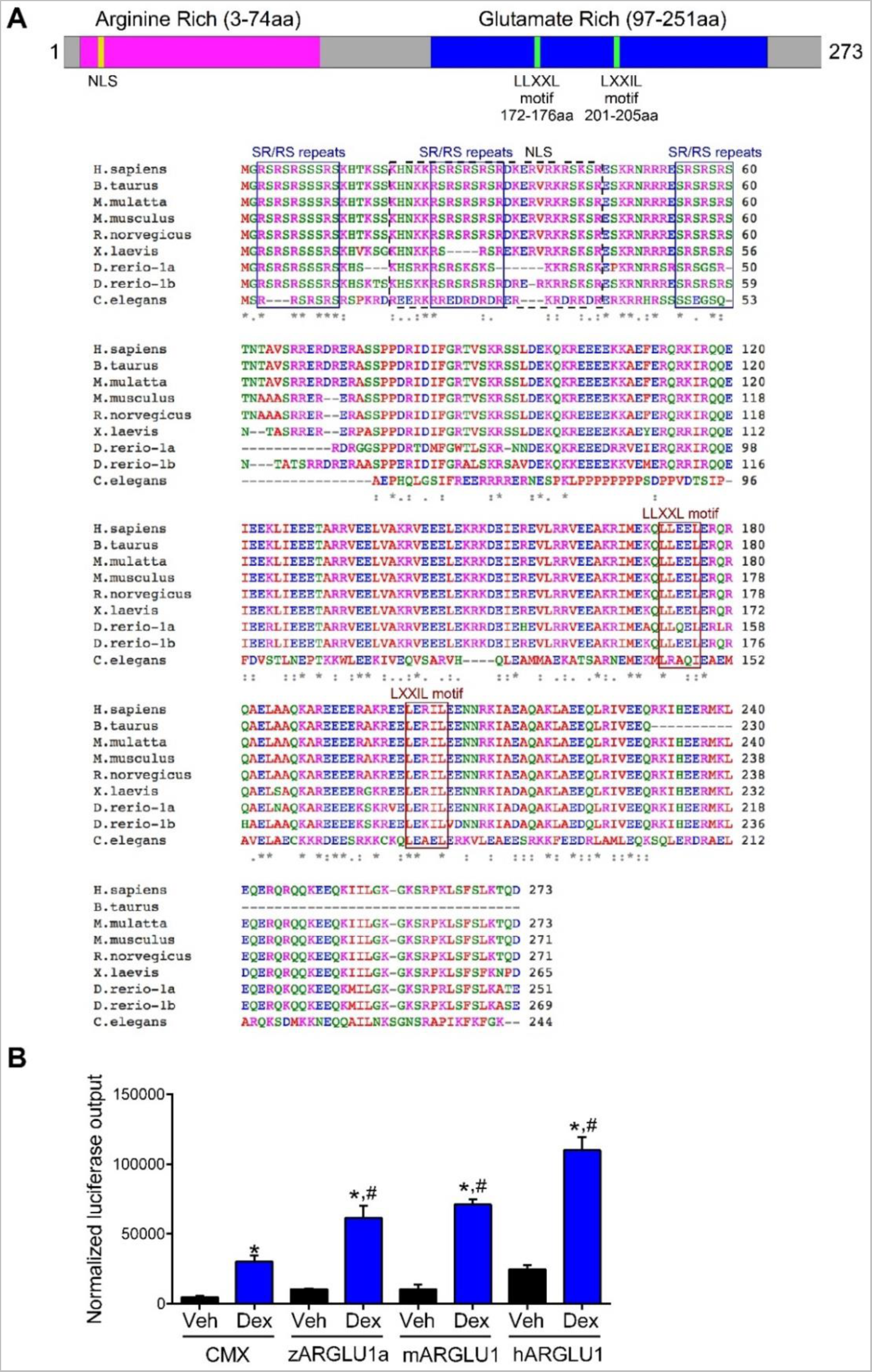
ARGLU1 is a highly evolutionary conserved GR coactivator, related to Figure 1. (**A**) Amino acid sequence alignment between species. ARGLU1 has two distinct regions: the N-terminus which is rich in positively charged arginine amino acids, and the C-terminus which is composed of glutamate rich amino acids. The arginine-rich region is also enriched in the SR/RS-repeats (boxed). Bipartite nuclear localization sequence (NLS) is depicted by a dashed box. Two putative NR interaction domains, LLXXL and LXXIL motifs, were identified by visual examination: L, leucine; I, isoleucine; X, any amino acid (boxed). Sequence alignment was performed using ClustalW. (**B**) ARGLU1's GR coactivation function is preserved across species. HEK293 cells were co-transfected with GAL4-GR and UAS-luciferase together with 15 ng of zebrafish (zARGLU1a), mouse (mARGLU1) or human (hARGLU1) ARGLU1. Six hours post-transfection, cells were treated with 100 nM dexamethasone (Dex) and harvested for luciferase assay 16 hrs later. β-galactosidase was used to normalize for transfection efficiency. When compared to the CMX group all three ARGLU1 orthologues were able to significantly induce GAL4-GR activity in response to Dex. Normalized luciferase output=luciferase light units/β-galactosidase*time. Data represent the Avg ± SD. *p<0.05 Dex group vs respective Veh; #p<0.05 vs CMX-Dex. ANOVA followed by Newman-Keuls test.

**Figure S2:**
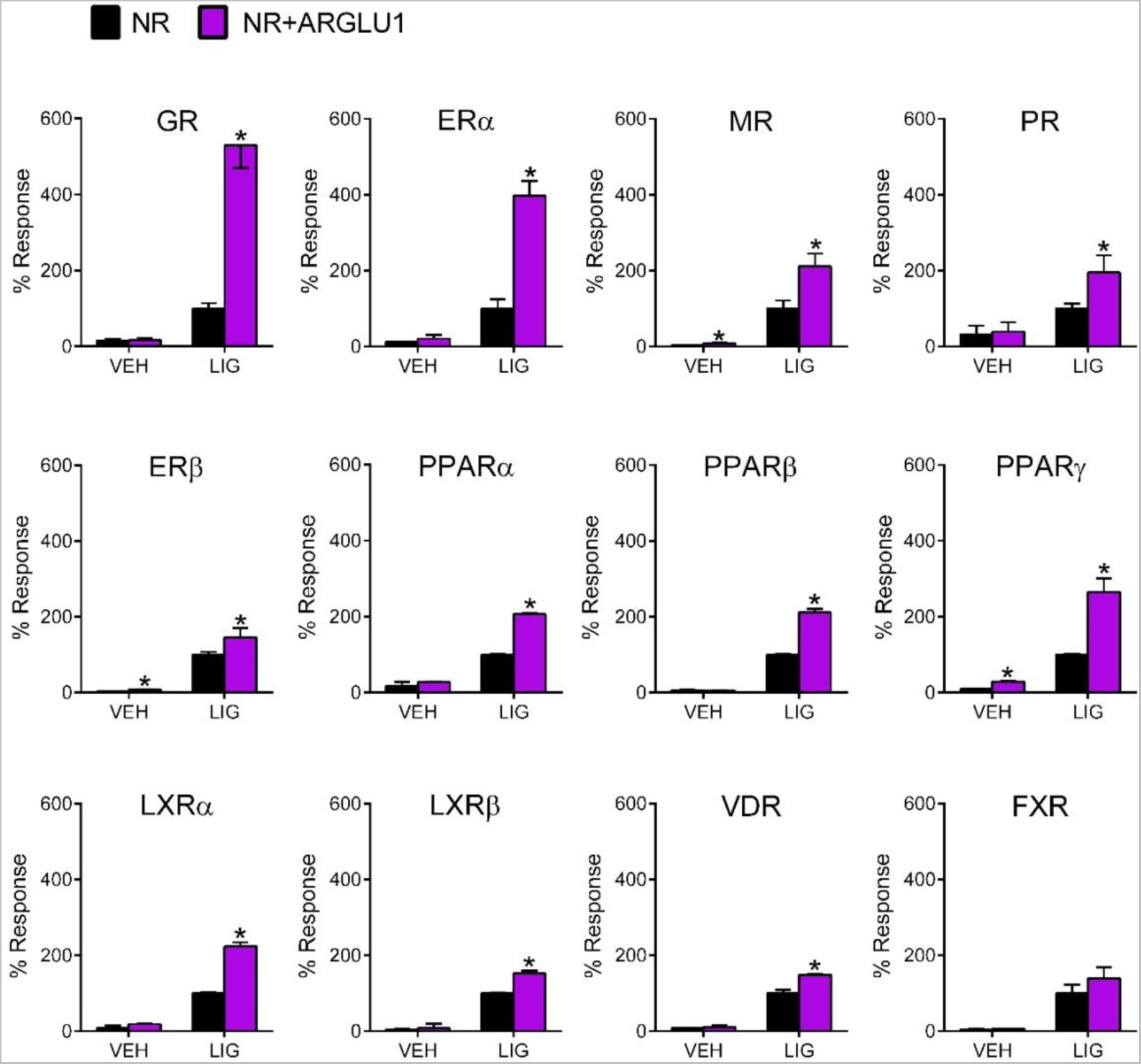
ARGLU1 activation of nuclear receptors, related to Figure 1. Various GAL4-nuclear receptor (NR) fusion proteins were co-transfected with the UAS-luciferase system and 15 ng ARGLU1 into HEK293 cells. Six hours later, cells were treated with 300 nM cortisol (hGRα), 10 nM aldosterone (hMR), 10 nM progesterone (hPR), 1 nM 17β-estradiol (hERα and hERβ), 500 nM T0901317 (hLXRα/hLXRβ), 500 nM WY14643 for (hPPARα), 25 nM GW1516 (hPPARδ), 50 nM rosiglitazone (hPPARγ), 10 μM CDCA (hFXR) and 0.5 nM 1,25(OH)_2_ vitamin D_3_ (hVDR). All ligands were dosed at the receptor’s EC_50_ in this system. Data represent the mean ± SD (n=3). *p<0.05 NR + ARGLU1 vs NR; Student’s t-test.

**Figure S3:**
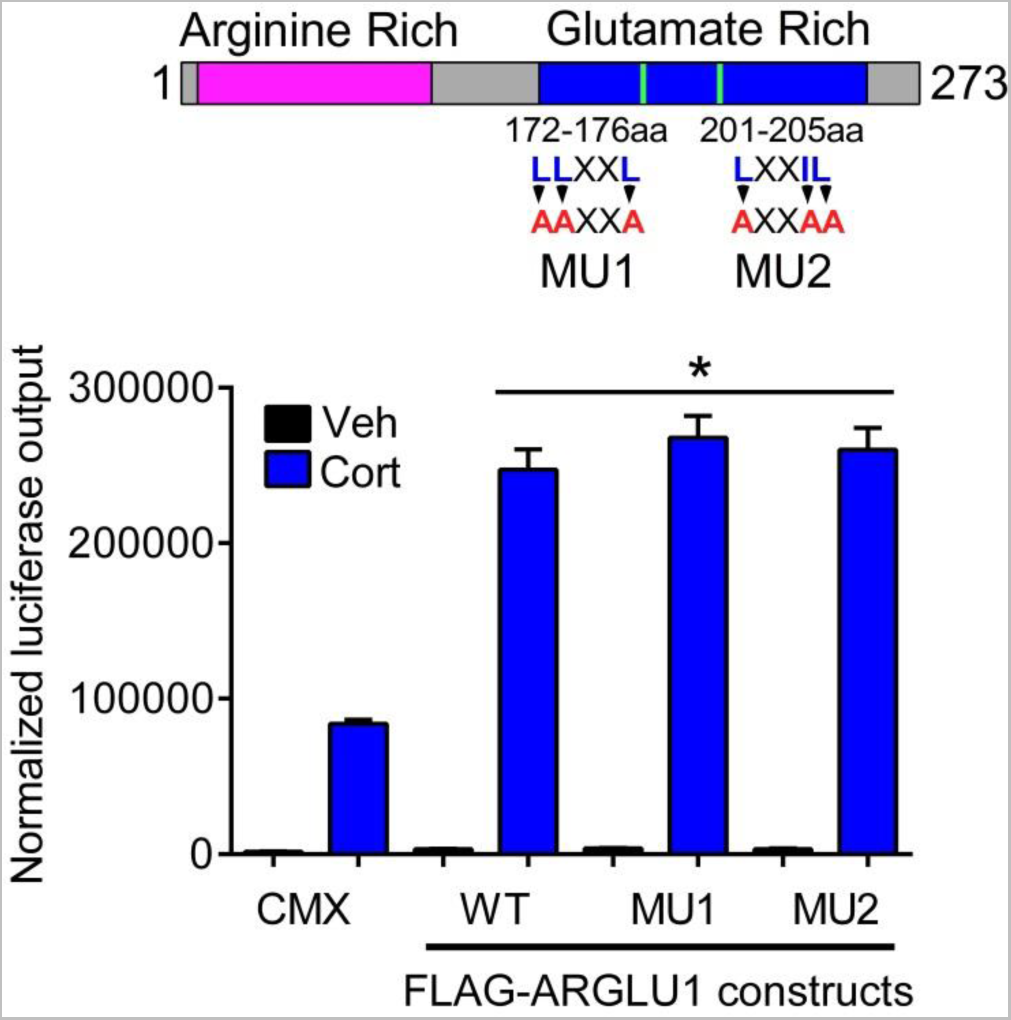
ARGLU1 activation of GR is not dependent on the individual LXXLL motifs, related to Figure 2. Top - Schematic diagram of ARGLU1 mutants. Bottom - GAL4-GR was cotransfected into HEK293 cells with the UAS-luciferase system and 15 ng of the indicated ARGLU1 LLXXL mutants, followed by administration of EtOH (Veh) or 300 nM cortisol. CMX was used as a control. Normalized luciferase output=luciferase light units/β-galactosidase*time. Data represent the mean ± SD (n=3). *p<0.05 vs CMX-Cort; ANOVA followed by Newman-Keuls test.

**Figure S4:**
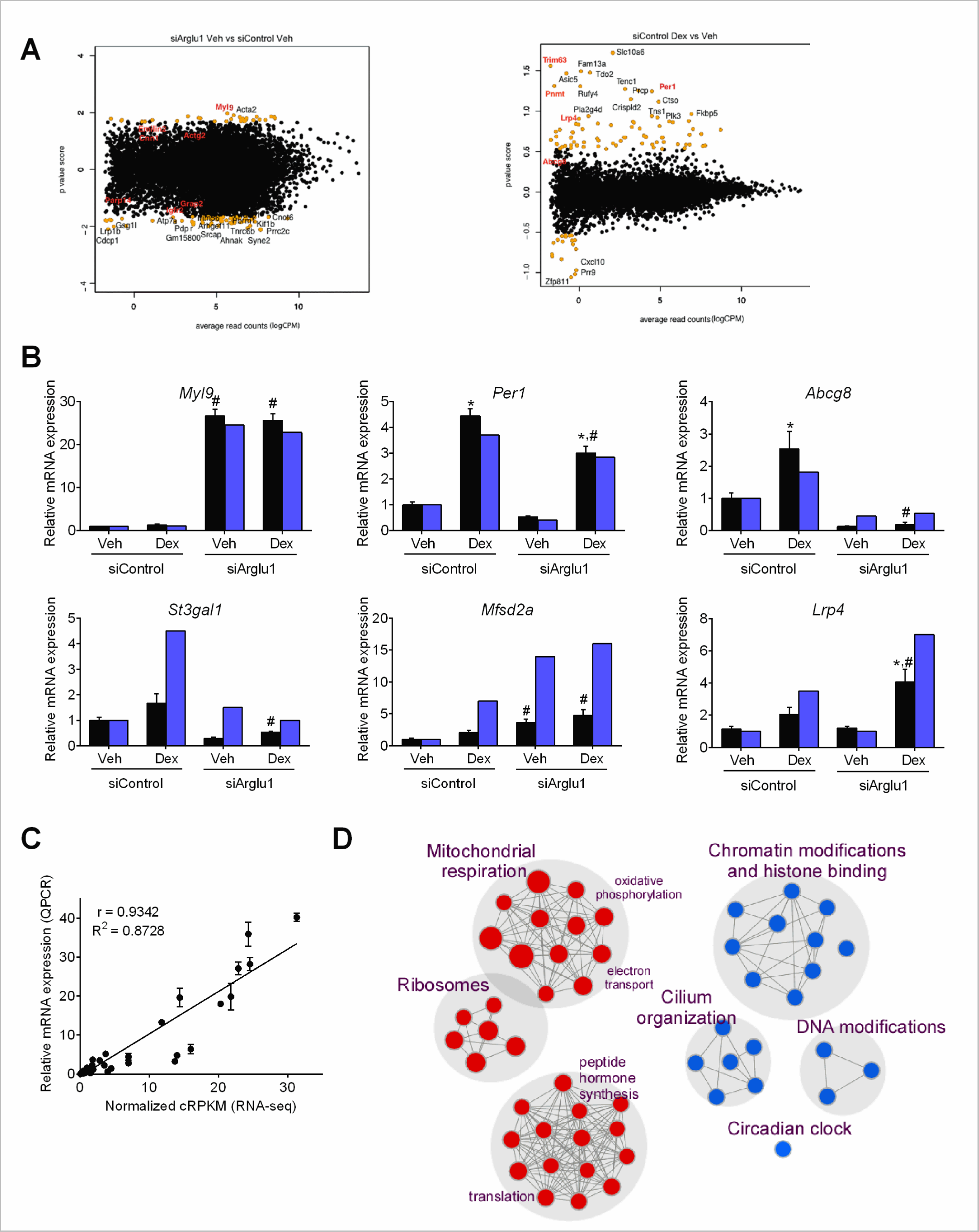
Loss of ARGLU1 in Neuro2a cells significantly alters gene expression in both GC-dependent and independent mechanisms, related to Figure 4. Neuro-2a cells were transfected with 30 pmol of siControl and siArglu1using RNAiMax for 48 hrs and then treated with vehicle (EtOH) or 100 nM Dex for 4 hrs before RNA extraction. (**A**) MA plot showing the distribution of read counts (x-axis) against p value score (defined in methods from edgeR analysis) for each gene. Orange - top 100 genes; Red – genes validated by QPCR. (**B**) QPCR validation of RNA-seq data was performed on non-pooled samples. Fold changes obtained from RNA-seq (cRPKM) are plotted for comparison. Data represent the mean ± SEM (n=3). *p<0.05 vs respective Veh, #p<0.05 vs siControl; ANOVA followed by Newman-Keuls test. (**C**) Correlation between fold change values from QPCR and RNA-seq data are shown without *Pnmt*. Inclusion of *Pnmt* yielded a correlation coefficient of r=0.9515. (**D**) GSEA analysis of basal changes in transcription upon ARGLU1 knockdown. Node size is proportional to the normalized enrichment score (i.e., the bigger the node size, the more significant this pathway was enriched). Red nodes –pathway upregulated by ARGLU1 knockdown. Blue nodes – pathway downregulated following ARGLU1 knockdown. Results are visualized using EnrichmentMap plug-in using the Cytoscape software.

**Figure S5:**
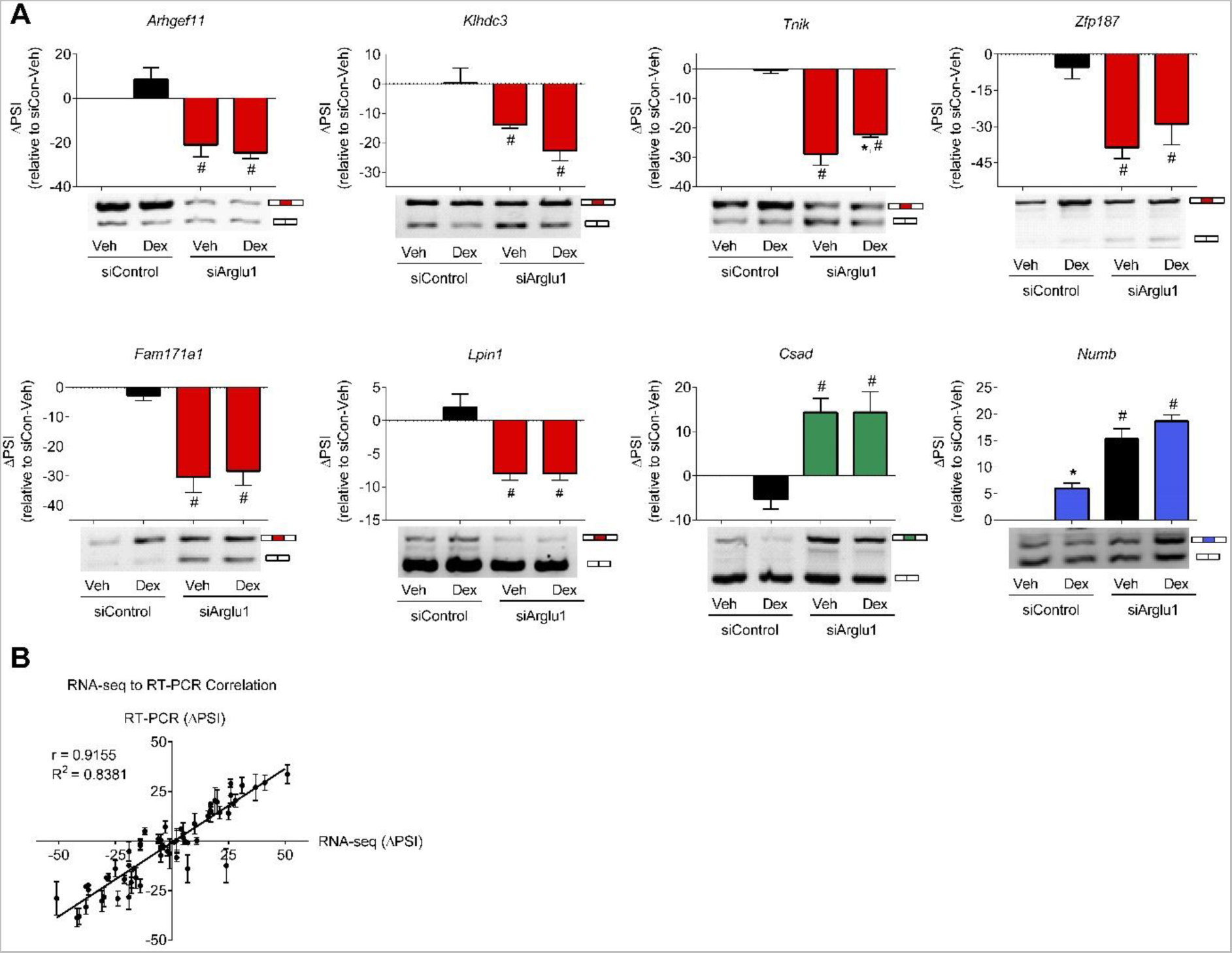
ARGLU1 knockdown leads to increased exon skipping and increased exon inclusion, related to Figure 5. (**A**) Splicing events with the PSI of ≥15 by RNA-seq were validated using one-step RT-PCR. Image J was used for quantification. PSI was calculated as: spliced in / (spliced in + spliced out) * 100%. ΔPSI is calculated by subtracting the individual PSIs from the PSI of siControl Veh group. Representative image is shown below the Image J quantification of the blot. Data represent the mean ± SEM (n=3). *p<0.05 vs respective Veh, #p<0.05 vs respective siControl; ANOVA followed by Neuman-Keuls test. (**B**) Correlation of ΔPSI values between RNA-seq and RT-PCR.

**Figure S6:**
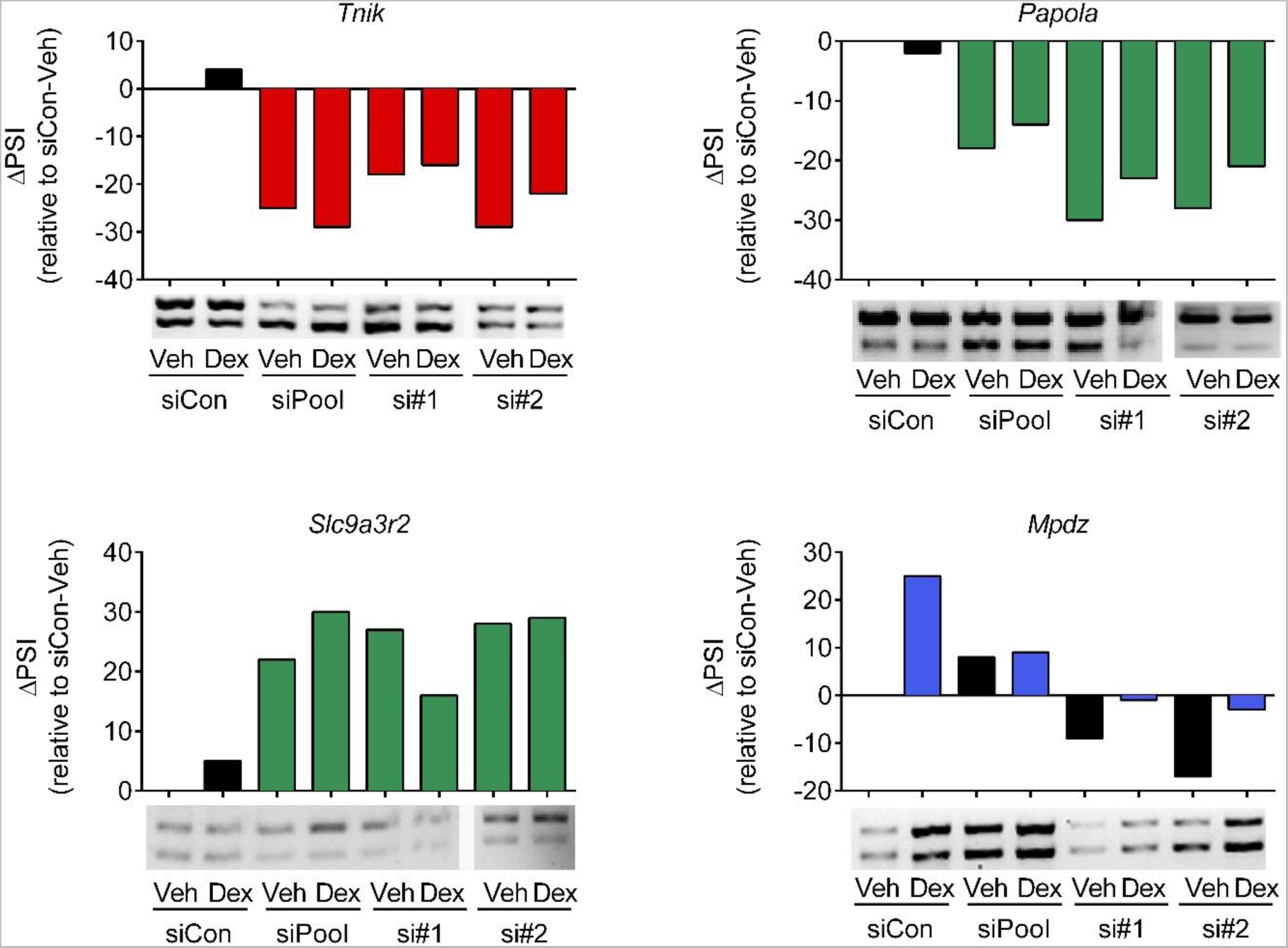
Changes in AS were confirmed with independent ARGLU1 siRNAs, related to Figure 5. N2a cells were transfected with 30 pmol of siControl or siRNAs targeting ARGLU1 (pool of 4 siRNAs, si#1 and si#2) using Lipofectamine RNAiMax reagent for 48 hrs and then treated with ethanol (Veh) or 100 nM Dex. a 4 hr ligand treatment RNA was extracted and one-step QPCR was used to analyze the indicated splicing events with a ΔPSI of ≥15 as identified by RNA-seq. siRNA#2 was used in the RNA-seq analysis. Image J was used for quantification. PSI was calculated as: spliced in / (spliced in + spliced out) * 100%. ΔPSI is calculated by subtracting the individual PSIs from the PSI of siControl Veh group. A representative image is shown below the Image J quantification of the blot.

**Figure S7:**
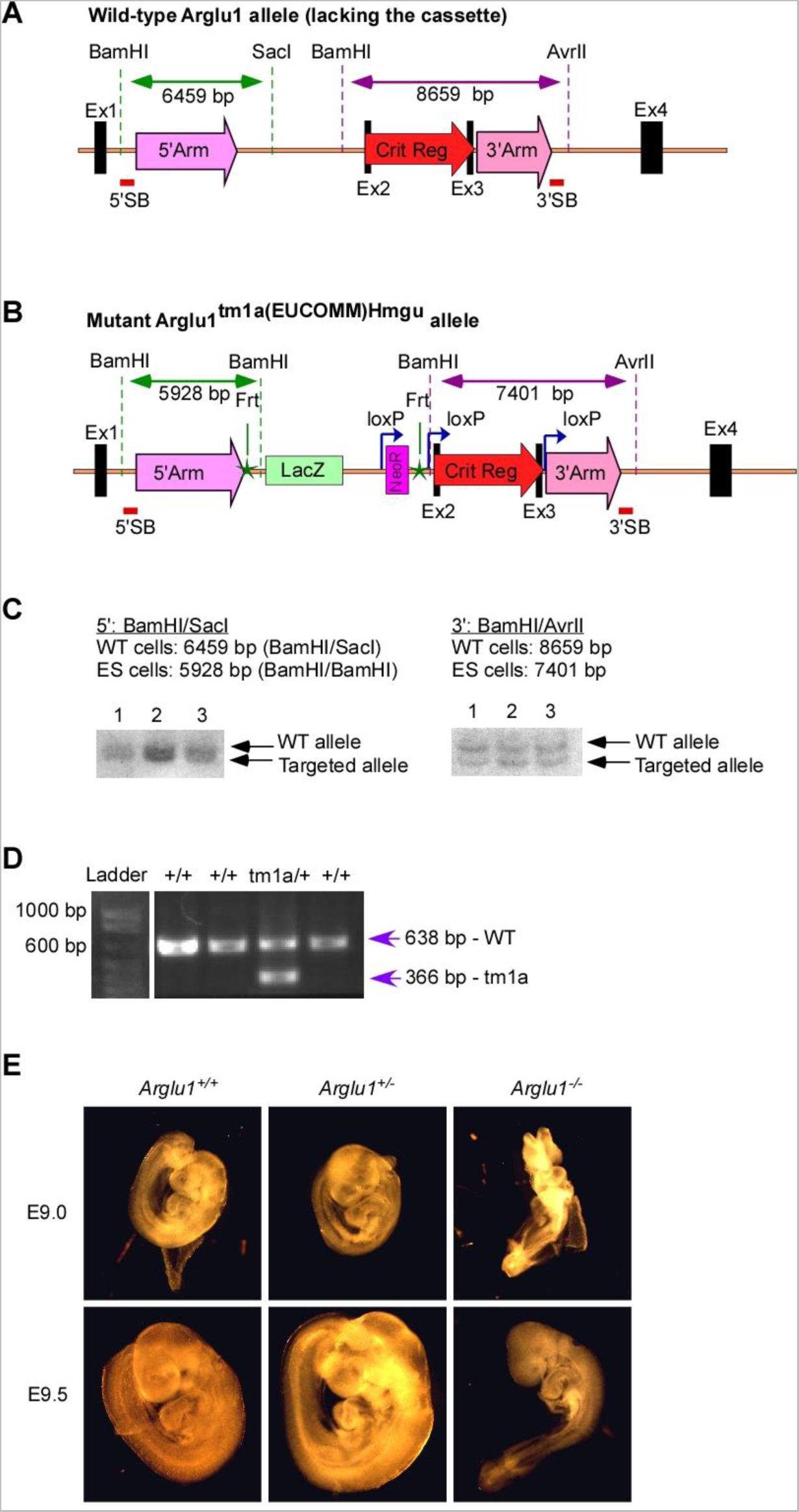
ES cell targeting construct and confirmation of germline transmission of ARGLU1^tm1a(EUCOMM)Hmgu^, related to Figure 7. Southern blot was used to confirm proper cassette targeting of the 3 different ES clones purchased from EUCOMM. (**A-B**) Schematic representation of the WT (**A**) and mutant (**B**) alleles. (**C**) Southern blot of digested ES clones. Critical region between exon 2 and 3 is targeted for deletion following Cre-recombinase generating Arglu1 null allele. All three clones confirmed proper cassette targeting. 5’ and 3’ Southern blot probes (5’SB and 3’SB, respectively) were generated using primers listed in extended experimental methods. To confirm the germ-line transmission (GLT) of the knockout first Arglu1^tm1a^ allele, male founder animals derived from the ES morula aggregation were crossed to albino female mice. After crossing to albino female mice, white pups were terminated at birth, non-white pups with black eyes were genotyped for tm1a germline transmission (**D**). (**E**) Mouse embryos from the *Arglu1^+/-^* x *Arglu1^+/-^* parent crosses were dissected and visually examined at E9.0 and E9.5. The yolk sac was used for genotyping. *Arglu1^+/-^* embryos appeared phenotypically normal whereas *Arglu1^-/-^* embryos had an overall developmental delay by approximately 0.5 days.

## Supplemental Material

### Supplemental Experimental Procedures

#### Cell lines and culture conditions

HEK293 and Neuro-2a (N2a) cells were maintained in high-glucose Dulbecco's modified Eagle medium (Sigma, St. Louis, MO) supplemented with 10% fetal bovine serum (Gibco, Carlsbad, CA) and 1 x penicillin/streptomycin (Sigma) at 37°C in a 5% CO_2_ humidified incubator. HEK293 cell transfections were performed in media containing 10% charcoal-stripped fetal bovine serum (Sigma) using calcium phosphate method in 96-well plates (Makishima et al., 1999). The total amount of plasmid DNA (150 ng/well) included 50 ng of luciferase reporter, 20 ng β-galactosidase, 15 ng nuclear receptor, varying concentrations of the nuclear receptor coactivators and pGEM filler plasmid. For co-immunoprecipitation experiments, HEK293 cells were grown in 100 mm plates and transfected with 5 µg of the indicated plasmids using calcium phosphate method. Neuro-2a (N2a) cells were transfected in suspension with siRNA against *Arglu1* (D-057082-02; 5’-GCCAAACGCAUCAUGGAAA-3’) or with the non-targeting Control siRNA (siGENOME Non-Targeting siRNA Pool #2; D-001206-14-05) using the RNAiMax reagent at a ratio of 1:6 (µL of RNAiMAX reagent to pmol of siRNA) to a final concentration of 12 nM of siRNA/well. Forty eight hours later, cells were treated with vehicle (ethanol) or 100 nM Dex in the growth media. Following a 4 hr ligand treatment, cells were harvested for RNA-seq and protein analyses.

#### High-throughput expression cloning screen

Electromax^TM^ DH10B^TM^ E.Coli cells (Invitrogen, Carlsbad, CA) were transformed with 10 ng of normalized human brain cDNA library (a gift from Dr. Stephane Angers). An aliquot (7.3 µL) of the transformation mixture was diluted into 3.2 L of Luria-Bertani broth supplemented with ampicillin (50 μg/mL). The diluted culture was then distributed into a deep 96-well plate (Sigma) at 1.4 mL/well (∼50 cDNA clones) and incubated for 27 hrs at 37 °C. Plasmid DNA was isolated from the bacterial cultures using the GenElute^TM^ HP 96 well plasmid midiprep kit (Sigma). HEK293 cells were maintained in high-glucose Dulbecco's modified Eagle's medium (Invitrogen) supplemented with 10% FBS (37 °C in 5% CO_2_). For transfection experiments, cells were seeded into clear bottom white 96-well plates (40,000 cells/well) in DMEM supplemented with 10% charcoal-stripped FBS. The following day, calcium phosphate method was used to transfect the DNA. The total amount of plasmid DNA (150 ng/well) included 50 ng UAS-luc reporter, 15 ng β-galactosidase, 15 ng of GAL4-hGR and 75 ng of cDNA library pool (50 clones/pool). For control reactions, 15 ng of pBK2-rTIF2 plasmid or 15 ng of CMX-RIP140 plasmid with 60 ng of CMX empty plasmid were used instead of cDNA pools. Six hours post-transfection, cells were treated with vehicle (ethanol) or 300 nM cortisol. Cells were harvested 14–16 hrs later for luciferase and β-galactosidase activity, as previously described (Makishima et al., 1999). Luciferase values were normalized to β-galactosidase to control for transfection efficiency and expressed as normalized luciferase output=luciferase light units/β-galactosidase*time. Data from triplicate wells were averaged and expressed as Avg ± SD.

#### Plasmid generation

The plasmids pIRES-puro-FLAG, pIRES-puro-mCherry, pIRES-puro-GFP and pcDNA5-FRT/TO-BirA* were gifts from Dr. Stephane Angers (University of Toronto, Toronto, Canada). GAL4-hGRα, CMX-hSRC1, CMX-hRIP140, CMX-mGR, UAS-luc, MMTV-luc, pGEM, β-galactosidase and CMX control plasmids were kindly provided by Dr. David Mangelsdorf (University of Texas Southwestern Medical Center, Dallas, TX). pBK2-rTIF2, GAL4-MR and PGC1α were gifts from Dr. Jason Matthews (University of Toronto). pBS SK(+)-islet-1 plasmid was a gift from Brian Ciruna (University of Toronto). pCMV-HA plasmid was purchased from Clontech. pCMV-SPORT6-hARGLU1, pCMV-SPORT6-mARGLU1, pExpress1-zARGLU1a, pME18S-FL-zARGLU1b, pCMV-SPORT6-hJMJD6, pCMV-SPORT6-hPUF60 and pOTB7-hU2AF2 plasmids were purchased from Open Biosystems. To generate FLAG-tagged hJMJD6, U2AF2 and PUF60 constructs, the respective coding regions were PCR-amplified from the plasmid DNA and sub-cloned into the pIRES-puro-FLAG vector using AcsI/NotI restriction sites. Full length human ARGLU1 and/or the indicated truncation mutants were PCR-amplified from a pCMV-SPORT6-hARGLU1 plasmid and subcloned into the CMX or pIRES-puro-FLAG, pIRES-puro-mCherry, or pcDNA5 FRT/TO plasmids using the KpnI/NheI or AscI/NotI restriction sites, respectively. Human GR was subcloned into pIRES-puro-FLAG plasmid from CMX-hGR plasmid with the AscI/NotI restriction sites. To generate HA-hGR and HA-hARGLU1 plasmids, GR and ARGLU1 were subcloned from CMX-hGR and CMX-hARGLU1 plasmids, respectively, with the KpnI/NotI sites into the pCMV-HA vector. The GST-expression plasmids for RNAcompete assays were constructed by inserting the coding region of full length ARGLU1, ARGLU1^N-term^ and ARGLU1^C-term^ into pTH6838 vector using AscI and SbfI restriction sites. Mutagenesis of the FLAG-ARGLU1 LXXLL-like motifs was performed using the QuikChange PCR mutagenesis kit (Agilent). Primers used for cloning the above constructs are listed below.

**Primer sequences used for plasmid generation.** All of the constructs were subcloned C-terminal to the tag.

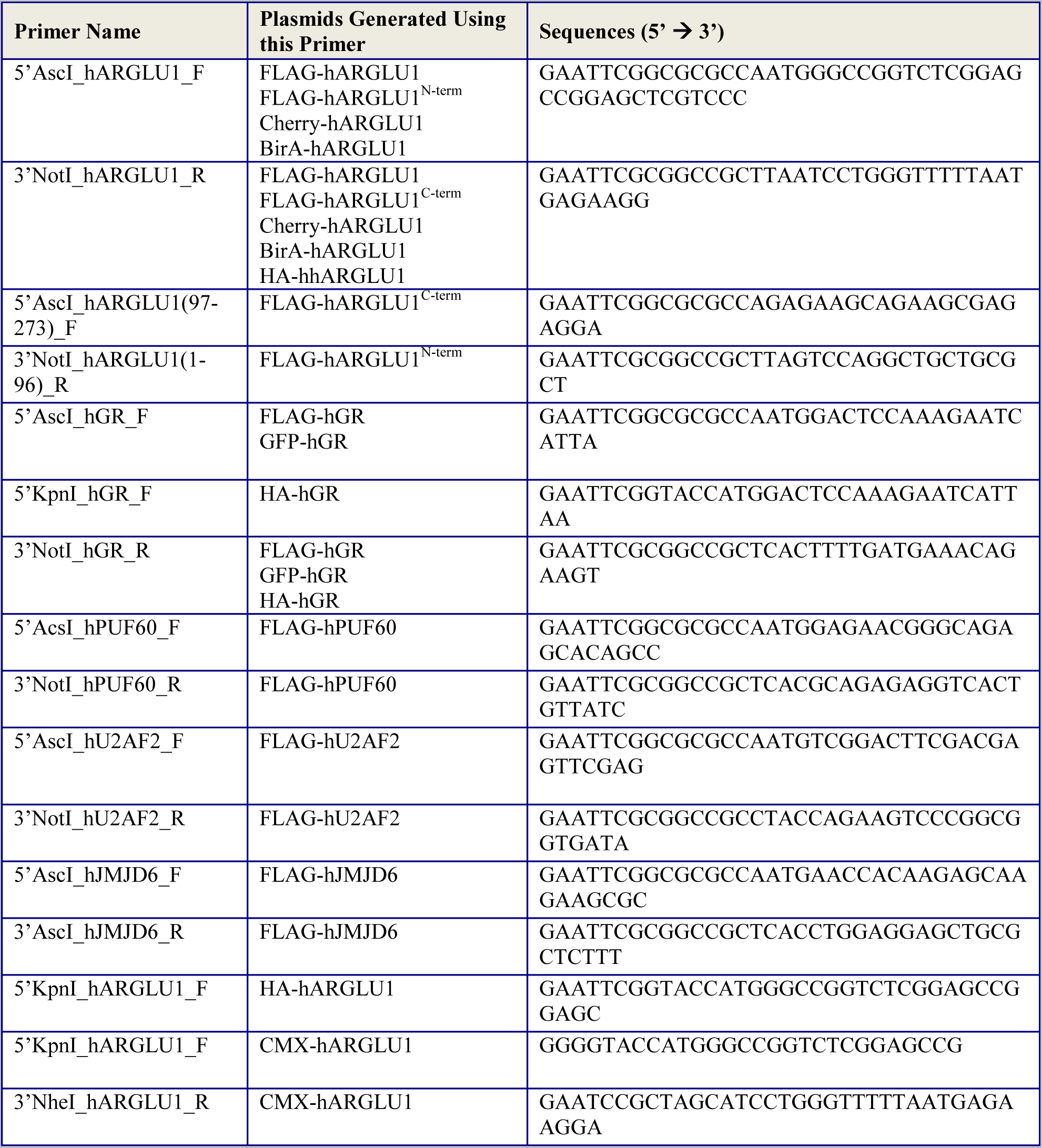
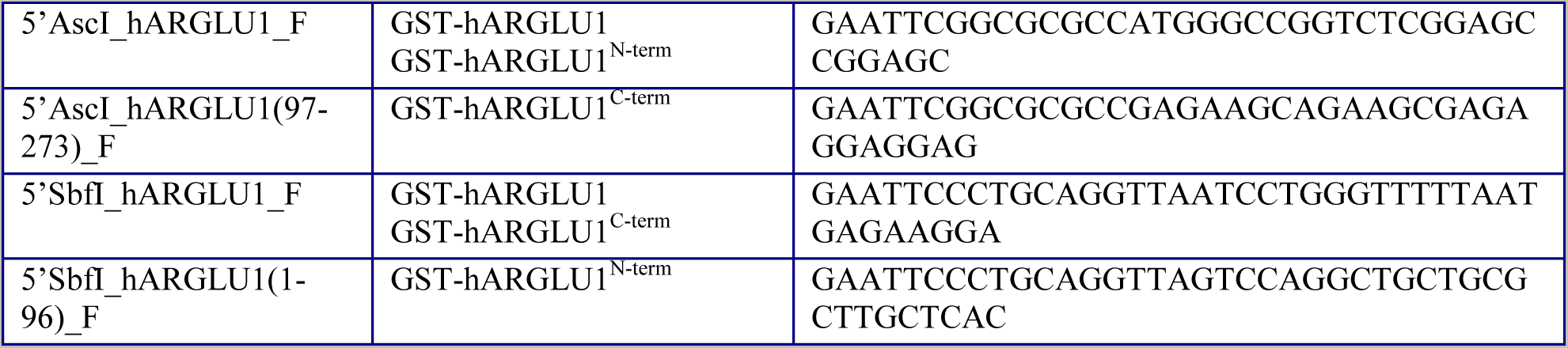

**Mutagenesis primers.**

Underlined are the mutated nucleotides relative to control.

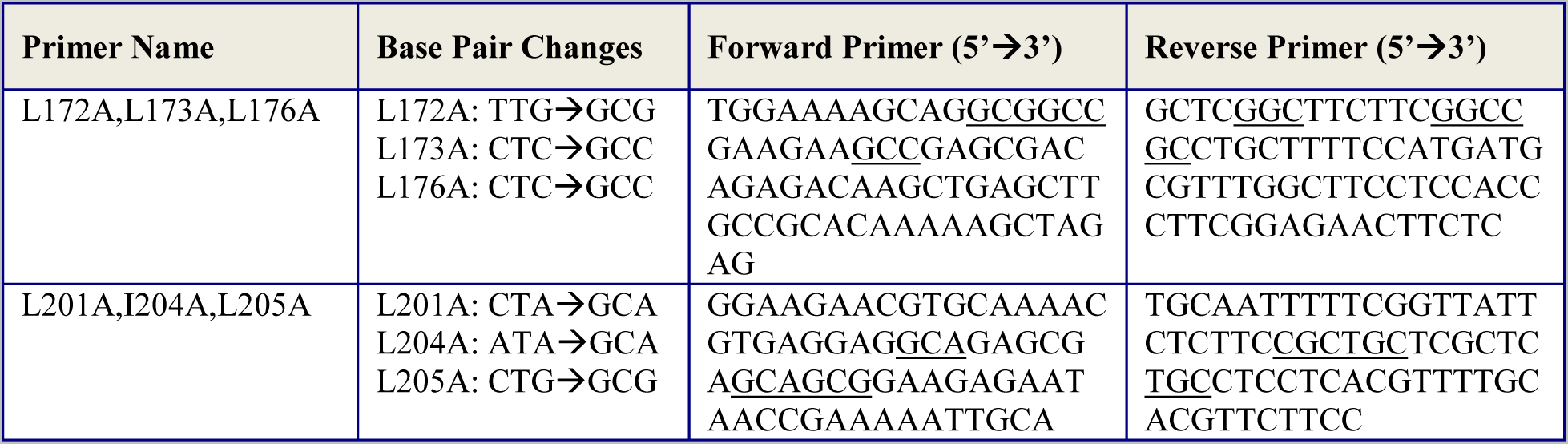

#### Tissue library generation

Four month old C57BL/6 (n=2-3) mice were sacrificed at lights on by isoflurane inhalation followed by exsanguination via the vena cava. Tissues were collected and immediately snap frozen in liquid N_2,_ except for pancreas which was processed immediately due to the high content of RNAse. Dorsal skin was shaved prior to collection. The digestive tract was sectioned, flushed with PBS and scraped mucosa was collected. White adipose tissue was obtained from the epididymal (eWAT) or inguinal (iWAT) area. Muscle-free bone was flushed with PBS to remove bone marrow. To obtain the spinal cord, the upper spinal column was sectioned (2 vertebrae/section) and the spinal cord was pushed out using a blunt tip of the forceps.

Tissues were stored at −80°C until RNA extraction. Ct values of ≥36 were considered undetectable for QPCR analysis.

#### Sequence analyses

NLS-MAPPER (Kosugi et al., 2009) [http://nls-mapper.iab.keio.ac.jp/cgi-bin/NLS_Mapper_form.cgi] was used to predict the nuclear localization sequence (NLS) in the ARGLU1 protein. ARGLU1 amino acid sequences from *Homo sapiens* (NP_060481.3), *Mus musculus* (NP_789819.2), *Rattus norvegicus* (NP_001020143.1), *Xenopus laevis* (NP_001090066.1), *Danio rerio* (1a-NP_998381.2 and 1b-NP_956456.1), *Caenorhabditis elegans* (NP_498950.1), *Bos taurus* (NP_001033279.1) and *Macaca mulatta* (NP_001180951.1) were aligned using ClustalW alignment program (Larkin et al., 2007) [http://www.genome.jp/tools/clustalw/].

#### Protein extraction

Cells were washed on the plate twice with ice-cold PBS, scraped, centrifuged at 700 x *g* for 5 min and lysed in 5 x the pellet volume of lysis buffer (50 mM Tris-HCl (pH 7.4), 1% Nonidet P-40, 150 mM NaCl, 1 mM EDTA, 1 mM EGTA, 0.1% SDS, 0.5% sodium deoxycholate (added fresh), 0.5 mM PMSF and 1 x protease inhibitors (Roche, Laval, QC, Canada) on ice for 20 minutes with occasional agitation. The lysates were centrifuged at 21,000 x *g* for 15 minutes at 4°C and the supernatants were transferred into a new tube and stored at -80°C until analysis. Protein concentration was assessed using the BCA assay (Pierce; Rockford, IL).

#### Western Blot analysis

Protein extracts (30 µg) were electrophoresed on a 4-20% gradient SDS gel and proteins were transferred in solution to a PVDF membrane (Bio-Rad, Hercules, CA). Membranes were blocked for 1 hour in 5% dried milk in TBS-T (50 mM Tris-HCl, pH 7.4, 150 mM NaCl, and 0.1% Tween 20) and incubated overnight at 4°C with primary antibodies: anti-ARGLU1 (1:600; Novus Biologicals, NBP1-87921), anti-βactin (1:1000; Cedarlane, ab8227), anti-HA (1:1000; Cell Signaling, 3724S) or anti-FLAG (1:1000; Sigma, F1804) in 1% milk in TBS-T (50 mM Tris-HCl, pH 7.4, 150 mM NaCl, and 0.1% Tween 20). The membrane was washed three times with TBS-T and incubated with secondary HRP-conjugated anti-rabbit IgG antibody (1:2000; Cell Signaling, 7074) or HRP-conjugated anti-mouse IgG antibody (1:5000; Cell Signaling, 7076) for 1 hr, followed by three more TBS-T washes. Samples were visualized with ECL-Prime and X-ray film (GE health care; Piscataway, NJ). The blots were quantified using Image J software.

#### Stable cell line generation

Stable HEK293 cells expressing FLAG-hARGLU1 were generated by transfecting 10 µg of pIRES-puro-FLAG-hARGLU1plasmid into 100 mm plate using the standard calcium phosphate transfection protocol (Makishima et al., 1999) followed by a subsequent selection with 2 µg/mL puromycin (Bioshop, Burlington, ON, Canada). *Flp***-**In™ *T-REx*™ 2*93* cells stably expressing BirA*-hARGLU1 were generated using the Flp-In system (Invitrogen) b*y* transfecting 1 µg of pcDNA5/FRT/TO-BirA-ARGLU1 plasmid together with 9 µg of pOG44 plasmid into 100 mm plate using a calcium phosphate transfection protocol (Makishima et al., 1999). Cells were then selected with 200 µg/mL Hygromycin B (Bioshop) in DMEM + 10% FBS.

#### FLAG affinity purification

Five 150 mm plates of HEK293 cells stably expressing FLAG-hARGLU1 were incubated with 500 nM Dex for 24 hrs, scraped, washed 2 x with ice-cold PBS and pelleted by centrifugation 700 x *g* for 5 min at 4°C. Pellets were snap frozen in liquid N_2_ and stored at -80°C until analysis. Cells were solubilized in 10 mL of TAP lysis buffer (50 mM HEPES-NaOH (pH 8.0), 100 mM KCl, 2 mM EDTA, 0.1% NP40, 10% glycerol, 1 mM PMSF, 1 mM DTT, and complete protease inhibitor cocktail (Roche)), and incubated for 1 hr at 4°C on a rotator. The protein lysates were centrifuged at 21,000 x *g* for 30 min at 4°C. The supernatant was transferred into a new tube and 50 μL of pre-equilibrated anti-FLAG M2 beads (A2220, Sigma) were added and incubated at 4 °C overnight. Beads were collected by centrifugation (700 x *g*, 1 min, 4°C), washed 3 x with TAP lysis buffer and 3 x with 50 mM ammonium bicarbonate pH 7.8. The proteins were eluted from the beads with 500 mM ammonium hydroxide (pH 11.0) in 3 × 100 µL fractions at 37°C. The eluates were transferred into a new centrifuge tube and lyophilized. Samples were resuspended in 100 µL of water, lyophilized again, resuspended in 50 mM ammonium bicarbonate and tryptic digest was performed overnight at 37°C with TPCK-trypsin (Promega, Madison, WI). Digested samples were lyophilized and resuspended in 45 µL buffer A (95% H_2_O, 5% acetonitrile, 0.1% formic acid). Next, 20 µL of the tryptic mixture was injected for LC-MS/MS analysis (LTQ-XL linear ion trap mass spectrometer, Thermo Scientific) as previously described (Zhang et al., 2014).

#### BioID assay

Five 100 mm plates (∼ 70% confluency) of *Flp***-**In™ *T-REx*™ 293 cells stably expressing BirA*-hARGLU1 were incubated for 24 hrs in complete growth media supplemented with 1 µg/mL tetracycline (Sigma), 50 µM biotin (BioShop) and 500 nM Dex. Cells were collected (700 x *g*, 5 min, 4°C), washed twice with ice-cold PBS, snap frozen in liquid N_2_ and stored at -80°C until analysis. Pellets were then lysed in 10 mL of modified RIPA lysis buffer (50 mM Tris-HCl pH 7.5, 150 mM NaCl, 1 mM EDTA, 1 mM EGTA, 1% Triton X-100, 0.1% SDS, complete protease inhibitors (Roche)) rotating at 4°C for 1 hr.

Lysates were sonicated 3 × 10 sec at a power of ∼20W (Misonix tip sonicator). Cell debris was pelleted by centrifugation at 21,000 x *g* for 30 min at 4°C. Clarified supernatants were incubated with 30 µL of packed, pre-equilibrated streptavidin-sepharose beads (GE 17-5113-01) at 4°C overnight. Beads were collected by centrifugation (700 x *g*, 1 min, 4°C), washed 2 x with modified RIPA lysis buffer, 2 x with TAP lysis buffer (50 mM HEPES-NaOH (pH 8.0), 100 mM KCl, 2 mM EDTA, 0.1% NP40, 10% glycerol, 1 mM PMSF, 1 mM DTT with complete protease inhibitor cocktail) and 5 x with 50 mM ammonium bicarbonate pH 8.3. Next, on-bead tryptic digest was performed with TPCK-trypsin (Promega) for 16 hrs at 37°C and the peptides were analyzed by mass spectrometry as described above.

#### Co-immunoprecipitation assays

HEK293 cells were transiently transfected with 5 µg of the expression plasmid of interest and treated with 500 nM Dex after 24 hrs. Forty eight hours later, cells were washed once with ice-cold PBS, pelleted by centrifugation 700 x *g* for 5 min at 4°C and washed twice more with ice-cold PBS. Pellets were snap frozen in liquid N_2_ and stored at -80 °C until analysis. Cells were solubilized in 1 mL of TAP lysis buffer (50 mM HEPES-NaOH (pH 8.0), 100 mM KCl, 2 mM EDTA, 0.1% NP40, 10% glycerol, 1 mM PMSF, 1 mM DTT, and complete protease inhibitor cocktail (Roche)), and incubated for 30 min at 4°C on a rotator. The protein lysates were centrifuged at 21,000 x *g* for 30 min at 4°C. The supernatant was transferred into a new tube and 30 μL protein G agarose beads were used to pre-clear the lysate. After pre-clearing, the lysate was split into 3 tubes and incubated at 4°C overnight with 5 µg of either HA, FLAG or IgG antibodies, as indicated. The next day, 50 μL of protein G agarose was added to the lysates and incubated on a rotator for 3-4 hrs at 4°C. Beads were collected by centrifugation (700 x *g*, 1 min, 4°C), washed 3 x with TAP lysis buffer and proteins were eluted in LDS containing β-mercaptoethanol.

#### RNA-seq studies and analysis

N2a cells were transfected with 30 pmol of siRNA targeting *Arglu1* and 48 hrs later mRNA was extracted as described above. Vehicle (ethanol) or 100 nM Dex was added to cells 4 hrs before harvest. mRNA was pooled by treatment group (n=3/group) and mRNA enriched Illumina TruSeq V2 RNA libraries were prepared. Samples were sequenced at the Donnelly Sequencing Centre (University of Toronto, Toronto, ON, Canada) on Illumina HiSeq2500 (averaging ∼196 million 100-nt paired-end reads per sample).

Transcriptome-wide AS and gene expression profiling were performed using a previously described workflow (Irimia et al., 2014). RNA-seq reads were aligned back to the mouse genome (Ensembl release 67) and transcript levels were quantified as reads per kilobase (of target gene) per million (of total reads) and corrected for transcript length and sequence redundancy (cRPKM) (Barbosa-Morais et al., 2012; Wang et al., 2009). cRPKM values were only used to correlate expression between RNA-seq and QPCR. The edgeR R Bioconductor package (3.12.0) was used on the gene read counts generated from the RNAseq aligned data to estimate differential expression between the treatment groups. Genes used as edgeR input were selected using a coverage threshold per gene that was set so that at least one sample had a count per million value (CPM + 1) ≥0.05. Dispersion parameters were estimated in edgeR by considering the pooled samples as replicates since each sample followed the same distribution. Proper data normalization and differential expression was assessed by MA plots (Figure S4A). The RNA-seq analysis including differential and exploratory plots (Volcano and MA plots) was performed using R/2.14.2. Changes in gene expression were validated by standard QPCR techniques (Bookout et al., 2006) using primers listed in the table below.

For alternative splicing analysis, Bowtie was used to align unique mappable positions for each junction. Percent spliced in (PSI) for each internal exon was defined as:

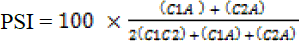 where C1A, C2A and C1C2 refer to the normalized read counts for each junction. Exons were considered “alternative” if the sample had 5 ≤ PSI ≤ 95. Splicing with quality scores of “LOW” or better were considered for analysis. Alternatively spliced genes which showed a change in PSI of 15 or more were validated using Qiagen One-Step RT-PCR kit. Primers were designed to flank the alternative exon (see below). Pooled mRNA was DNase-treated and One-Step RT-PCR was carried out as per the manufacturer's instructions except the reactions were performed in a 10 µL volume. Briefly, a master mix was made to include 0.4 mM dNTPs, 600 nM of forward and reverse primers and the Qiagen enzyme mix. A total of 1 ng of DNAse-treated RNA was used in each reaction. Cycle times and annealing temperatures are listed for each of the primer sets listed below. Splice products were resolved on a 2% agarose gel. Image J was used to quantify band intensity for each spliced variant (referred to as in or out) and then the PSI metric was calculated as follows: PSI=spliced in / (spliced in + spliced out) * 100%.

**Primers used to validate gene expression using QPCR**

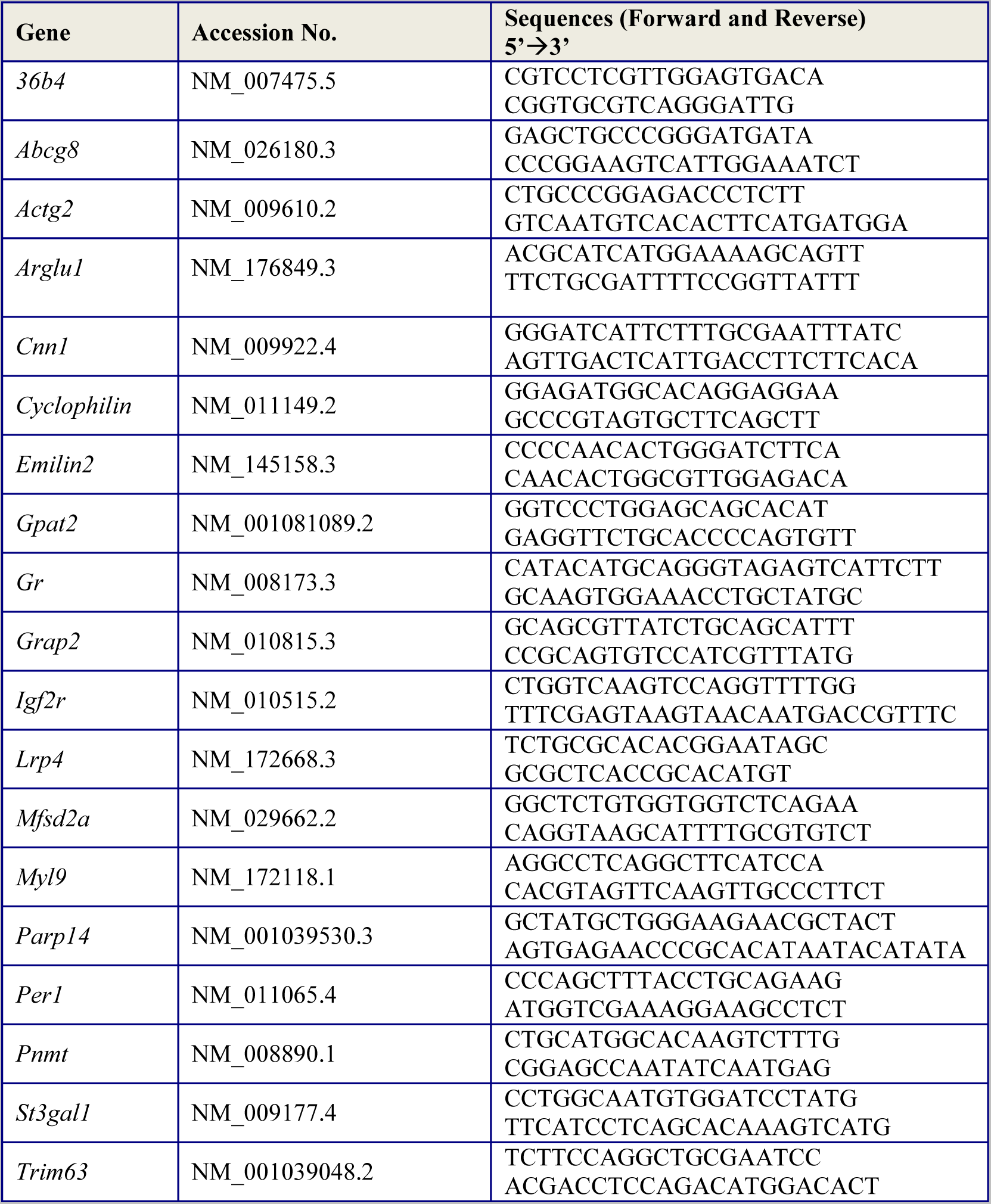

**Primers used to validate alternatively spliced genes using one-step RT-PCR**

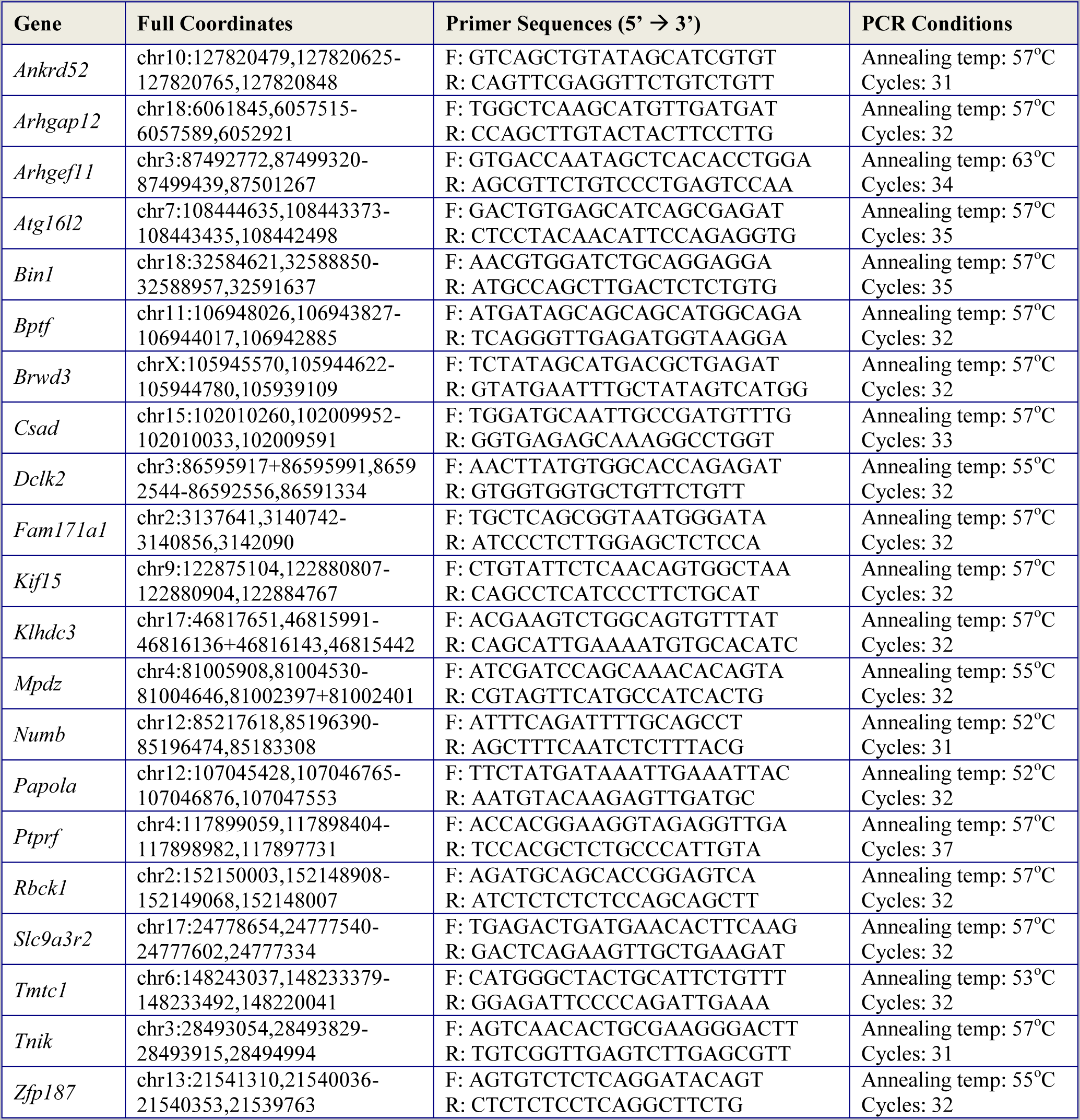

#### RNA Immunoprecipitation studies

Three 10 cm plates of N2a cells were grown until 60-70% confluency before transfection. Cells were transfected with FLAG-CMX, FLAG-hARGLU1, FLAG-hARGLU1(1-96), or FLAG-hARGLU1 (97-273) using Lipofectamine^TM^ 3000 reagent (Invitrogen) following the usual protocol from the manufacturer. Cells were collected 48 hrs after transfection and protein-RNA complexes were crosslinked using formaldehyde to a final concentration of 1% for 10 mins then quenched by glycine to a final concentration of 125 mM. Cells were collected, pelleted by centrifugation and washed twice with ice-cold PBS. The cell pellet was resuspended with 3 volumes of swelling buffer (5 mM Hepes [pH 8], 85 mM KCl, 0.5% NP40 and protease inhibitor cocktail) and incubated on ice. Nuclei were pelleted by centrifugation at 2,500x *g* for 5 mins. Nuclei were resuspended in nuclei lysis solution (50 mM Tris-HCl [pH 8.1], 10 mM EDTA [pH 8], 1% SDS (w/v), protease inhibitor cocktail and RNase inhibitor (ThermoScientific) and incubated on ice. Sample was diluted in FA lysis buffer (1 mM EDTA [pH 8], 50 mM HEPES-KOH [pH 7.5], 140 mM NaCl, 0.1% SDS, 1% TritonX-100 and PIs) and sonicated using a Misonix on the highest power output (10W) for 5 rounds with 1 min on/2 min off cycles. Extracts were cleared by centrifugation for 10 mins at 4°C and transferred to new tubes. Initial DNase treatments of the extracts were performed, then samples were centrifuged and transferred to new tubes. 10% input RNA was taken and flash frozen in liquid nitrogen. 20 µl bead volume (50 µl of 50% slurry) of FLAG M2 magnetic beads (Sigma) were added to the extracts and incubated with end-over-end rotation overnight at 4°C. Sequential 5 min washes were performed with FA lysis buffer, FA500 buffer, LiCl wash buffer, and TE buffer. Two elutions were performed at 37°C while mixing at 3,000 rpm using RIP elution buffer. Proteins were digested with proteinase K at 45°C for 45 mins and then crosslinking was reversed with NaCl at 70°C for 45 mins. RNA was extracted using RNA-STAT60 and the RNA pellet was resuspended in DEPC-H_2_O. ARGLU1 binding to RNA was assessed using specific qPCR primers crossing intron-exon boundaries to look at pre-mRNA binding. One-Step RT-qPCR (Froggabio kit) was performed on a 7900HT machine and quantitated using dilutions of the 10% input for the standard curve (ABI Biosystems).

#### Pathway enrichment analysis

To run gene set enrichment analysis using the GSEA software (http://software.broadinstitute.org/gsea/index.jsp), a ranking score was calculated by applying the formula 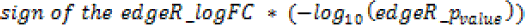 which ranks genes from most up-regulated to most down-regulated for each comparison between two treatment groups. This ranked list of genes was imported into the GSEA software and gene set enrichment analysis was performed using the pre-ranked option and 2000 gene set permutations, a minimum gene set size of 5 and a maximum gene set size of 500 as parameters. The gene sets included in the GSEA analyses were biological pathways obtained from KEGG, MsigDB-c2, NCI, Biocarta, IOB, Netpath, HumanCyc, Reactome and the Gene Ontology (GO) databases, updated August 2014 (http://baderlab.org/GeneSets). EnrichmentMap is a useful tool which organizes large gene data sets, such as pathways and gene ontology terms, into a network where closely related gene sets are clustered together, allowing for an easier visualization and interpretation of the data. EnrichmentMap (version 2.1, Cytoscape 3.2.1) was used to visualize enriched gene-sets with a nominal p-value<0.01 and a Jaccard overlap coefficient of 0.35 (Figure S4D).

To generate a gene expression and splicing overlap map (Figure 6B), gene set enrichment analysis was performed using the g:Profiler software (Reimand et al., 2007). For splicing, 928 genes with dPSI of ≥15 were analyzed, whereas for the analysis of the gene expression data set, genes with a p-value<0.05 (607 genes) were uploaded into g:Profiler. g:Profiler parameters were set to have a size of functional category between 3 and 500, and size of Q&T (size of the overlap) to 3. The gene sets included in the analyses were obtained from KEGG, GO Biological Process and Reactome.

#### Generation of ARGLU1 knockout animals

Three targeted ES cell lines (HEPD0639_4_A04, HEPD0639_4_B06 and HEPD0639_4_C03) heterozygous for the knockout-first-reporter tagged *Arglu1^tm1a(EUCOMM)Hmgu^* allele (Figure S7A,B) were obtained from European Conditional Mouse Mutagenesis Program (EUCOMM). ES cells were expanded at the Toronto Centre for Phenogenomics (TCP) and proper cassette targeting was confirmed using Southern blot analysis (Figure S7C). Chimeric founder animals were generated at the TCP facility by morula aggregation of two individual ES clones (HEPD0639_4_A04 and HEPD0639_4_B06). Animals with more than 50% chimerism (13 in total) were set up for germline-test breeding with albino female mice obtained from Charles River (B6N-*Tyr^c-Brd^*/BrdCrCrl; strain code #493). Black eyed pups with non-white coat color were then tested for germline transmission of the Arglu1^tm1a^ allele by PCR of the tail genomic DNA (Figure S7D). Arglu1 heterozygous null mice (*Arglu1*^+/-^) were then bred together in an attempt to generate homozygous null animals (*Arglu1*^-/-^). Pregnant dams were sacrificed at E9.0, E9.5 and E12.5 days post coitus to determine when embryonic lethality occurred. The yolk sacs were used to genotype embryos. Genotyping primers are listed below.

#### Southern Blot analysis

Genomic DNA from 3 ES cell clones was prepared using standard phenol/chloroform extraction. Ten µg of DNA was digested with BamHI/SacI or BamHI/AvrII at 37°C overnight and separated on a 1% agarose gel. The DNA was denatured, transferred onto Nytran SPC membrane using a Whatman TurboBlotter kit (GE health care, Piscataway, NJ) and hybridized with the indicated ^32^P-labeled probes (RediPrime, Amersham, Buckinghamshire, UK). The 5’ and 3’ external probes, 151 bp and 328 bp in size, respectively, were generated by PCR from genomic DNA using primers listed below. Membranes were washed at 65°C twice with 2 x SSC (Sigma) with 0.2% sodium dodecyl sulfate (5 min/wash) and once with 0.2 x SSC with 0.1% sodium dodecyl sulfate for 20 min and exposed to X-ray film (Kodak) for 36 hrs. The BamHI/SacI Southern analysis yielded two bands of 6459 bp (wild-type allele) and 5928 bp in size (tm1a allele). The BamHI/AvrII Southern blot exhibited two distinct bands of 8659 bp (wild-type allele) and 7401 bp in size (tm1a allele).

**Southern Blot and genotyping primer list.**

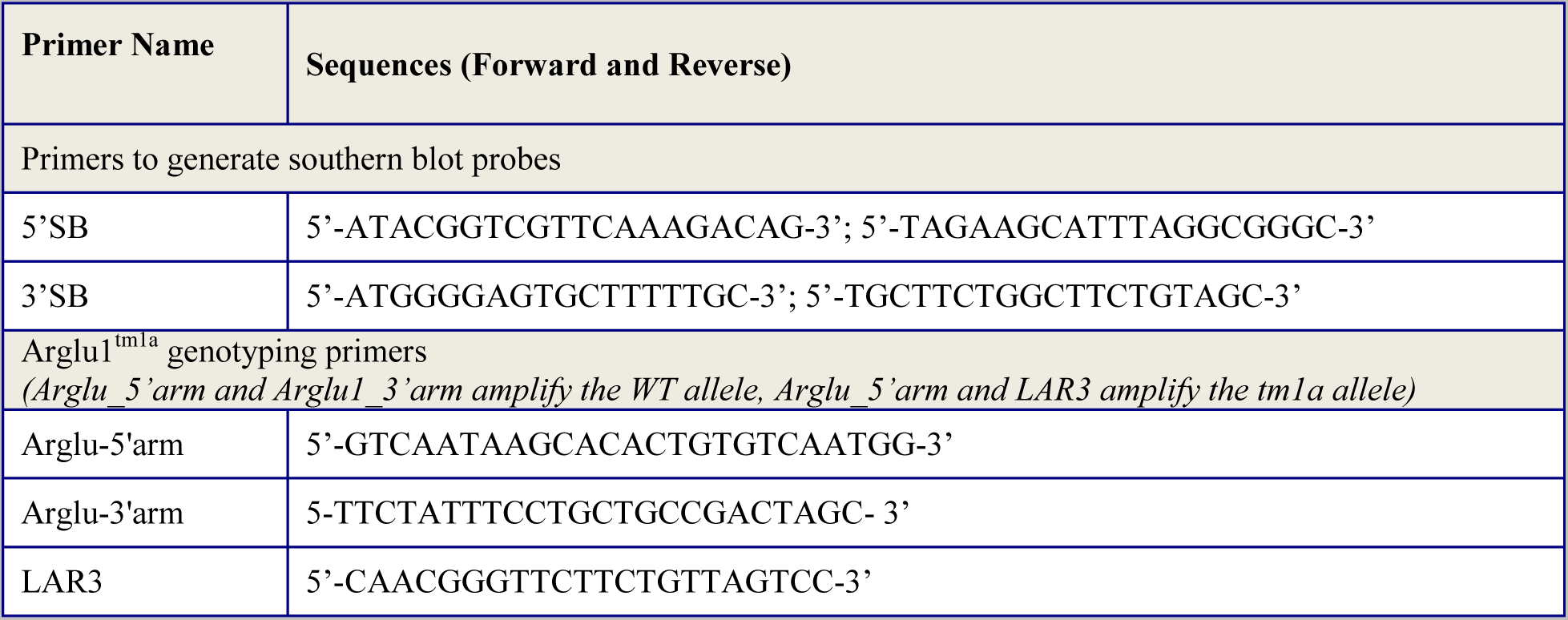

#### Zebrafish morpholino injections and rescue

Translation-blocking morpholinos (MO) targeting zebrafish *arglu1a*, *arglu1b* and *p53* genes were purchased from Gene Tools, LLC (Philomath, OR). Morpholino injections were carried out using standard techniques. 4.5 ng of p53 MO alone or in combination with 4.8 ng of arglu1a MO and/or 4.8 ng of arglu1b MO made in 0.1% phenol red was injected into the one-cell stage embryo. *Arglu1a* and *arglu1b* rescue RNA was synthesized by SP6 transcription from the PCR-based template using the mMESSAGE mMACHINE system (Ambion Inc, Austin, TX). PCR templates were amplified from plasmid DNA using Platinum High-Fidelity Taq-Polymerase (Invitrogen) under standard PCR conditions. The gene-specific forward primer contained an artificially introduced SP6-promoter to enable synthesis of the sense transcript (primer sequences are listed below). PCR fragments were gel-purified using Quick-spin columns (Valencia, CA). Purified template DNA was used for *in vitro* transcription. 0.1 ng of the indicated rescue RNA was injected into the embryos as above. Embryos were imaged at indicated times after fertilization.

#### Zebrafish whole mount *in situ* hybridization

*Arglu1a* and *arglu1b* antisense RNA probes (∼500 bp) were generated from a PCR-based template. PCR templates were amplified from plasmid DNA using Platinum High-Fidelity Taq-Polymerase (Invitrogen) under standard PCR conditions. The gene-specific reverse primer contained an artificially introduced T7-promoter to enable synthesis of the antisense transcript. PCR fragments were then gel-purified using Quick-spin columns (Qiagen, Valencia, CA). Purified template DNA was used for *in vitro* transcription incorporating DIG-UTP with the mMESSAGE mMACHINE system (Ambion; Austin, TX). Islet-1 probes were generated from gel-purified BamHI linearized pBS SK(+) plasmid using T3 polymerase. Labeled RNA probes were then purified by standard ammonium acetate purification. The concentration and quality of RNA probes were assessed by running a 1% TAE agarose gel. Whole mount *in situ* hybridization was carried out as previously described (Hauptmann and Gerster, 1994). Embryos were cleared in 100% methanol, mounted in benzylbenzoate:benzylalcohol (2:1) and images were taken with Leica M205 FA microscope.

**Primers used in zebrafish experiments.**

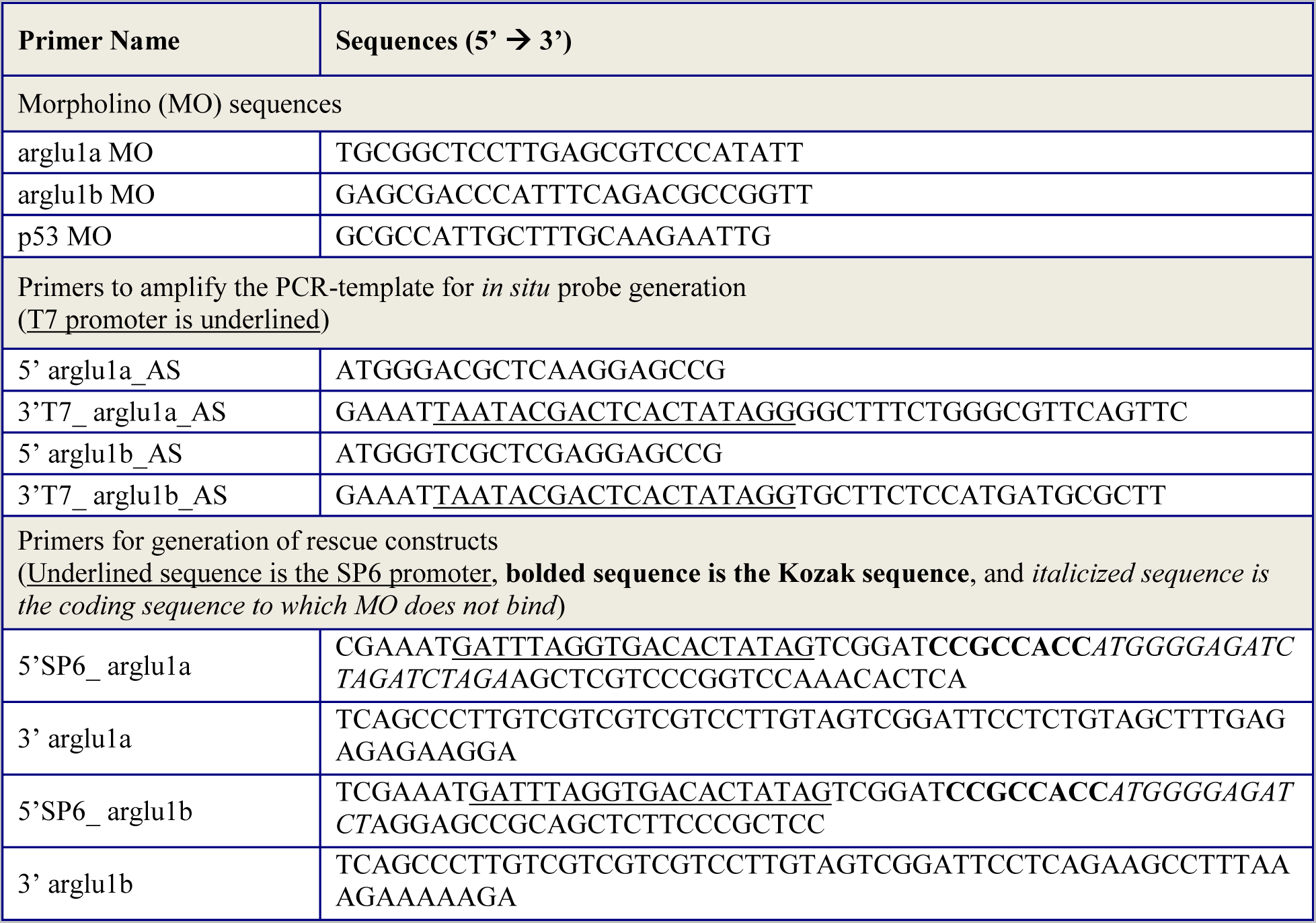

